# Optogenetic silencing of neurotransmitter release with a naturally occurring invertebrate rhodopsin

**DOI:** 10.1101/2021.02.18.431673

**Authors:** Mathias Mahn, Inbar Saraf-Sinik, Pritish Patil, Mauro Pulin, Eyal Bitton, Nikolaos Karalis, Felicitas Bruentgens, Shaked Palgi, Asaf Gat, Julien Dine, Jonas Wietek, Ido Davidi, Rivka Levy, Anna Litvin, Fangmin Zhou, Kathrin Sauter, Peter Soba, Dietmar Schmitz, Andreas Lüthi, Benjamin R. Rost, J. Simon Wiegert, Ofer Yizhar

**Author notes:** These authors contributed equally to this work (Mathias Mahn, Inbar Saraf-Sinik, Pritish Patil, Mauro Pulin).

## Abstract

Information is carried between brain regions through neurotransmitter release from axonal presynaptic terminals. Understanding the functional roles of defined neuronal projection pathways in cognitive and behavioral processes requires temporally precise manipulation of their activity *in vivo*. However, existing optogenetic tools have low efficacy and off-target effects when applied to presynaptic terminals, while chemogenetic tools are difficult to control in space and time. Here, we show that a targeting-enhanced mosquito homologue of the vertebrate encephalopsin (eOPN3) can effectively suppress synaptic transmission through the G_i/o_ signaling pathway. Brief illumination of presynaptic terminals expressing eOPN3 triggers a lasting suppression of synaptic output that recovers spontaneously within minutes *in vitro* as well as *in vivo*. In freely moving mice, eOPN3-mediated suppression of dopaminergic nigrostriatal afferents leads to an ipsiversive rotational bias. We conclude that eOPN3 can be used to selectively suppress neurotransmitter release at synaptic terminals with high spatiotemporal precision, opening new avenues for functional interrogation of long-range neuronal circuits *in vivo*.

## Introduction

Neurons typically form both local and long-range synaptic connections, through which they interact with neighboring neurons and with distant neuronal circuits. Long-range neuronal communication is crucial for synchronized activity across the brain and for the transmission of information between brain regions with distinct information processing capabilities. For example, dopaminergic neurons in the substantia nigra project to the dorsal striatum via the nigrostriatal pathway and play a critical role in movement control as part of the basal ganglia circuitry (Alcaro, et al., 2007). Manipulating the activity of such long-range projection pathways allows a detailed evaluation of their functional contribution to cognitive and behavioral processes and has become a widespread approach through the use of optogenetic and chemogenetic techniques. However, while optogenetics allows robust and temporally-precise excitation of long-range projecting axons (Yizhar, et al., 2011), silencing such long-range connections with existing optogenetic tools has proven difficult (Wiegert, et al., 2017). We have previously shown that the light-driven chloride pump halorhodopsin (eNpHR3.0) only partially suppresses neurotransmitter release. The proton-pumping archaerhodopsin (eArch3.0) triggers off-target effects, including an increase in intracellular pH and elevated spontaneous neurotransmission (Mahn, et al., 2016), potentially leading to off-target behavioral consequences (Lafferty & Britt, 2020). While halorhodopsin-mediated inhibition has no effect on intra-synaptic pH (Mahn, et al., 2016), it does temporarily shift the chloride reversal potential and can lead to GABA-mediated excitation (Raimondo, et al., 2012). Furthermore, both halorhodopsin and archaerhodopsin require continuous delivery of high light power to sustain their ion pumping activity (Zhang, et al., 2007). Alternative approaches, such as optogenetic induction of synaptic plasticity (Creed, et al., 2015; Klavir, et al., 2017; Nabavi, et al., 2014), or inhibition by disruption of the release machinery (InSynC (Liu, et al., 2019); photo-uncaging of botulinum toxin-B (Liu, et al., 2019)), can effectively decrease synaptic transmission, but are not as temporally precise as direct optogenetic manipulations.

Chemogenetic tools such as DREADDs (Designer Receptors Exclusively Activated by Designer Drugs) or PSAMs (Pharmacogenetically Selective Actuator Modules; (Armbruster, et al., 2007; Magnus, et al., 2011)) pose an alternative to optogenetics for manipulation of presynaptic terminals. It has been shown that presynaptic terminal function can be effectively suppressed by delivery of the cognate ligands of these engineered receptors (Basu, et al., 2016; Stachniak, et al., 2014). However, these approaches require direct infusion of the ligand to the location of the targeted presynaptic terminals, and their temporal specificity is fundamentally limited by the binding affinity to and clearance of the ligand. The DREADD hM4Di has been shown to act as an inhibitor of synaptic transmission (Stachniak, et al., 2014). By activating the G_i/o_ pathway, hM4Di suppresses the synaptic release machinery through a mechanism similar to that of endogenous presynaptic inhibitory GPCRs, presumably through suppression of Ca^2+^ channel activity (Herlitze, et al., 1996) and inhibition of the vesicle release machinery downstream of Ca^2+^ entry (Gerachshenko, et al., 2005; Zhu & Roth, 2014; Zurawski, et al., 2019). We reasoned that a light-activated G_i/o_-coupled rhodopsin could potentially trigger the same type of synaptic suppression (Fig. 1A). However, while many known vertebrate rhodopsins do couple to the G_i/o_ pathway, these proteins are difficult to utilize as optogenetic tools since they undergo photobleaching after G protein dissociation as part of their natural phototransduction cycle (Bailes, et al., 2012) (Fig. 1B). Previous studies have revealed that bistable type-II rhodopsins are abundant across vertebrates and invertebrates (Tsukamoto & Terakita, 2010). These photoreceptors form a stable association with both the cis- and trans-configuration of the retinal chromophore (similar to the microbial type-I rhodopsin family including channelrhodopsin) and are therefore often referred to as bistable photopigments (Koyanagi, et al., 2004; Terakita, 2005). Importantly, bistable type-II rhodopsins show reduced photobleaching (Bailes, et al., 2012) (Fig. 1B). We reasoned that members of the bistable type-II rhodopsin family that couple to G_i/o_ signaling would be suitable candidates for light-mediated silencing of neurotransmitter release from presynaptic terminals.

**Figure 1.**
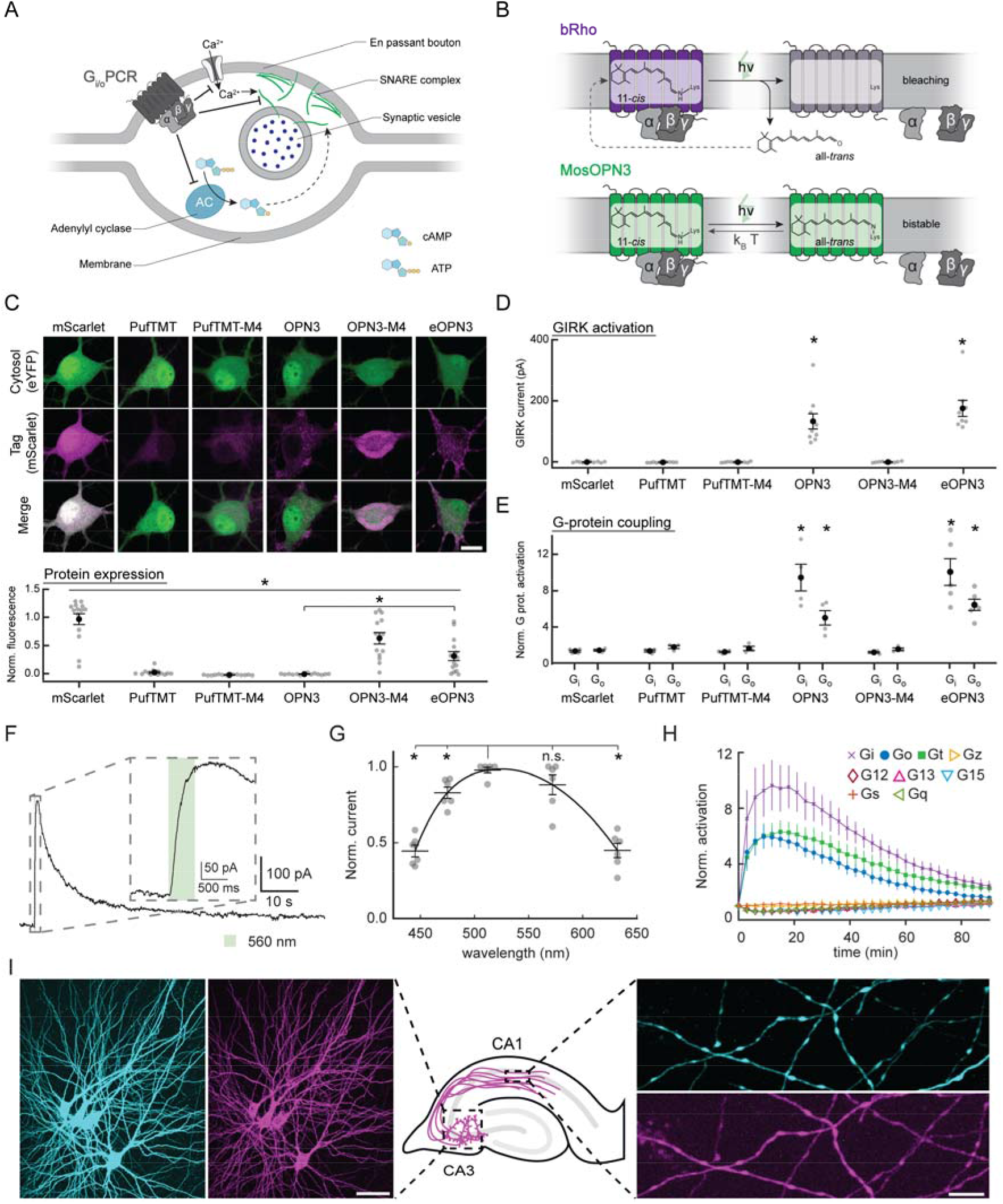
G_i/o_-coupled rhodopsins for light-mediated presynaptic inhibition. **(A)** Schematic diagram depicting the mechanism through which G_i/o_ signaling reduces the synaptic vesicle release probability. An activated GPCR leads to inhibition of voltage-gated Ca^2+^ channels as well as reduced cAMP levels, both leading directly (solid arrow) and indirectly (dotted arrow) to a reduction of Ca^2+^-dependent vesicle release. **(B)** Schematic diagram of distinct retinal binding mechanisms in bleaching (*top*) and bistable (*bottom*) rhodopsins. Bleaching rhodopsins release all-*trans*-retinal following photon absorption (*h*·*v*) and need to bind a new 11-*cis*-retinal before being able to enter the next photocycle. Bistable rhodopsins sustain their covalent bond with retinal independent of its configuration, removing the influence of 11-*cis*-retinal tissue availability. In bistable rhodopsins, all-*trans***-** retinal switches back to 11-*cis*-retinal either by absorbing another photon or spontaneously in the dark with a probability depending on the kinetic energy of the molecule (*k*_B_·T). *k*_B_ = Boltzmann constant; T = thermodynamic temperature; *h* = Planck constant; ν = photon frequency. (**C**) Representative confocal images of neurons co-transfected with expression vectors for eYFP and the indicated rhodopsin variants. Images show fluorescence in the eYFP channel (top), the mScarlet channel (middle) and the merged images (bottom). Bottom: Expression level of each of the displayed rhodopsin-mScarlet constructs, quantified as the average pixel intensity in n > 13 neurons for each construct normalized to cells expressing only mScarlet. The amount of measured fluorescence differed between all conditions (p = 1.34·10^−12^ Kruskal-Wallis test followed by Bonferroni-Holm corrected pairwise comparisons using Wilcoxon rank sum tests: OPN3 vs. eOPN3 fluorescence n = 14, p = 1.3·10^−4^. Scale bar, 15 μm. Images in the mScarlet channel are individually scaled for visualization of low fluorescence levels. Fluorescence measurements were taken under matched imaging conditions for all variants tested. (**D**) GIRK currents evoked by a 500 ms pulse of 560 nm light at 2 mW·mm^−2^ in GIRK2-1 co-expressing hippocampal neurons during a voltage clamp recording, held at −70 mV. Only cells expressing OPN3 and eOPN3 showed GIRK-mediated currents (p = 1.71·10^−6^ Kruskal-Wallis test followed by Bonferroni-Holm corrected pairwise comparisons using Wilcoxon rank sum tests). (**E**) Light-dependent G protein activation by opto-GPCR constructs assayed in HEK293T cells expressing an individual GsX chimera and a cAMP reporter. Only OPN3 and eOPN3 showed G_i_ and G_o_ activation. See fig. S2 for complete assay and statistics. **(F)** Sample whole-cell voltage-clamp recording of a cultured hippocampal neuron co-expressing eOPN3 and GIRK2-1, held at −70 mV. The depicted current was evoked by a 500 ms light pulse of 2 mW·mm^−2^ at 560 nm. Inset shows an expanded view of the GIRK current onset during the light pulse. (**G**) Action spectrum of endogenous GIRK-mediated currents in neurons expressing eOPN3, normalized to peak activation per cell (n = 6, p = 3.45·10^−4^ Friedman rank sum test followed by pairwise comparisons using Conover’s test). Peak excitation occurred at 512 nm (p < 4.24·10^−3^ Holm corrected pairwise comparisons to all other wavelengths except 572 nm). **(H)** Light-dependent G protein activation by eOPN3, assayed as in fig. S2. eOPN3 specifically and strongly activated inhibitory G proteins (G_i_, G_o_, G_t_) in a light-dependent manner (n = 5). See fig. S2 for complete assay and statistics. (**I)** eOPN3 expression in CA3 pyramidal neurons in organotypic hippocampal slice cultures. Two-photon maximum-intensity projections of CA3 neurons co-expressing the cytosolic fluorophore mCerulean (cyan) and eOPN3-mScarlet (magenta). Shown are the somatodendritic compartment of neurons electroporated with the two plasmids (*left; s*cale bar, 50 μm) and their axons projecting into *stratum radiatum* of CA1 (*right; s*cale bar, 5 μm). Plots depict individual data points and average ± SEM.

Here, we tested several bistable rhodopsin variants for use as optogenetic tools, specifically addressing their expression in mammalian neurons and their capacity for G_i/o_ pathway activation and light-driven inhibition of presynaptic release. While many of these invertebrate opsins failed to express in mammalian neurons, we were able to optimize the expression of a mosquito-derived homolog of the mammalian encephalopsin/panopsin protein (OPN3). The mosquito OPN3 is a bistable photopigment that allows high-efficiency and specific recruitment of the G_i/o_ signaling cascade (Koyanagi, et al., 2013). Using our targeting-enhanced OPN3 (eOPN3) protein, we were able to suppress synaptic release in rodent hippocampal, cortical and mesencephalic neurons. In behaving mice, eOPN3 triggered robust pathway-specific behavioral effects, suggesting that eOPN3, and potentially other members of the bistable rhodopsin family, can be utilized as optogenetic tools for potent silencing of the activity of presynaptic terminals with high spatiotemporal precision.

## Results

### Expression of naturally-occurring and engineered G_i/o_-coupled bistable rhodopsins in mammalian neurons

We reasoned that the efficient suppression of presynaptic function by the DREADD hM4Di (Fig. 1A, (Stachniak, et al., 2014)) arises from the stable binding of the engineered ligands of these receptors (Sternson & Roth, 2014) and the subsequent, stable G_i/o_-mediated signal transduction. We therefore hypothesized that rhodopsins coupling to the G_i/o_ pathway could serve as potent presynaptic silencing tools provided that persistent activation of such a tool can be achieved with light. While vertebrate visual rhodopsins, which dissociate from their retinal chromophore upon illumination (Fig. 1B, bRho), can in principle be used for presynaptic silencing (Li, et al., 2005), it remains unclear whether these rhodopsins can provide sufficiently robust activation of the G_i/o_ pathway at presynaptic terminals to support potent and sustained effects. Recent work has identified several new members of the encephalopsin subfamily of ciliary opsins, that couple to the G_i/o_ pathway. Encephalopsins exist in a wide range of organisms, including the pufferfish teleost multi-tissue opsin (PufTMT) and the mosquito OPN3 (OPN3). These rhodopsins are intrinsically bistable, as they retain the covalent bond between the retinal chromophore and the protein moiety (Fig. 1B) and display prolonged signal transduction following activation (Koyanagi, et al., 2013). We tested several photoreceptors of this family for expression in mammalian neurons. To maximally recapitulate the signaling pathway of the M4 acetylcholine receptor, as utilized by hM4Di, we also generated chimeric photoreceptors composed of bistable invertebrate rhodopsins and the intracellular domains of M4 (Fig. S1).

To evaluate the utility of these bistable rhodopsins and their M4 chimeras, we first characterized their expression and membrane targeting in neurons (Fig. 1C). We transfected primary cultured hippocampal neurons with mammalian codon-optimized versions of PufTMT, OPN3 and M4 chimeras of these rhodopsins, with C-terminal mScarlet fusions for direct visualization. Next, we measured their ability to evoke G protein-coupled inwardly-rectifying potassium channel-mediated (GIRK) currents in cultured neurons as a readout for functional activation of the G_i/o_ pathway. We conducted whole-cell recordings in neurons co-transfected with plasmids encoding one of each of the rhodopsin variants along with a GIRK2-1 channel (Lesage, et al., 1994). This configuration allowed us to quantify and compare the magnitude of G_i/o_ pathway activity through the measurement of GIRK2-1-mediated hyperpolarizing K^+^-currents. The endogenous expression of GIRK2-1 in neuronal cell types (Lüscher & Slesinger, 2010) and its ability to form functional homotetramers (Whorton & MacKinnon, 2011) make GIRK2-1 well suited as a reporter of G_i/o_ pathway activation in neurons. In parallel, we determined the interactions between the rhodopsin variants and specific G proteins using a HEK cell-based GPCR screening assay that couples the opsin to a G_s_-chimera (GsX assay, Fig. S2A, (Ballister, et al., 2018)). This approach allowed us to analyze their interaction with all major G proteins (G_i_, G_o_, G_t_, G_q_, G_s_, G_z_, G_12_, G_13_, G_15_), based on chimeras of the respective G-protein with G_s_, which triggers bioluminescence in a cAMP-dependent manner.

Both the wild-type PufTMT opsin and the PufTMT-M4 chimera displayed low levels of expression in cultured neurons and did not yield light-activated GIRK currents (Fig. 1C-D). Our GPCR screen showed that illumination of PufTMT-expressing cells only activated G_z_ (Fig. 1E; Fig. S2B). Koyanagi and colleagues previously demonstrated that PufTMT can recruit the G_i/o_ pathway only with highly concentrated PufTMT, nevertheless inducing efficient inhibition of cAMP production after illumination (Koyanagi, et al., 2013). Our results suggest that this adenylyl cyclase inhibition is mediated by G_z_ signaling. In combination, these results suggest that PufTMT cannot be used to fully recapitulate the efficient inhibition of vesicle release induced by hM4Di. In contrast with PufTMT, the wild-type mosquito OPN3 protein (referred to as OPN3 hereafter) yielded GIRK-mediated currents in all recorded neurons (Fig. 1D) and coupled efficiently with G_t_, G_i_ and G_o_ signaling cascades (Fig. 1E and S2B), in contrast to other rhodopsins (Spoida, et al., 2016). However, in mammalian neurons, the expression of OPN3 was low, punctate, and mostly intracellular (Fig. 1C). These results suggested that enhancing the membrane targeting of OPN3 will improve its efficacy as an optogenetic tool for synaptic terminal silencing. The OPN3-M4 chimera, containing the intracellular loops of the M4 acetylcholine receptor, expressed at higher levels in comparison to OPN3, but showed a predominantly intracellular localization (Fig. 1C). Moreover, OPN3-M4 did not evoke any detectable GIRK currents (Fig. 1D) or G protein activation (Fig. 1E; Fig. S2B).

### Generation and characterization of a targeting-enhanced OPN3

Our previous work indicated that addition of an ER export signal (ER) along with a Golgi trafficking signal (ts) to the light-gated chloride channel GtACR2 (eGtACR2; (Mahn, et al., 2018)) led to an increase in axonal membrane localization. We therefore modified OPN3 in a similar manner, yielding the enhanced OPN3-ts-mScarlet-ER (eOPN3). This modification led to an increased overall expression and enhanced membrane targeting (Fig. 1C) in cultured hippocampal neurons, compared to OPN3. Green light pulses delivered to neurons co-expressing eOPN3 and GIRK2-1 channels triggered robust GIRK-mediated currents (Fig. 1D, F). Activation of GIRK currents was maximal at 512 nm (Fig. 1G), consistent with previous characterization of light absorption by OPN3 protein (Koyanagi, et al., 2013).

We confirmed that eOPN3 retained its capacity to specifically activate the G_i/o_ pathway using the GsX assay. Light-activation of GsX-expressing HEK cells yielded selective and strong activation of G_i_-, G_o_- and G_t_-mediated signal transduction, but not of other G proteins (Fig. 1E, H and Fig. S2B). To rule out undesired consequences of heterologous rhodopsin overexpression, such as impaired cell health or light-independent effects on the physiological activity of expressing neurons, we examined the intrinsic excitability of cultured hippocampal neurons expressing eOPN3-mScarlet. Whole-cell patch-clamp recordings revealed no significant difference in intrinsic properties between neurons expressing eOPN3-mScarlet and neighboring, non-expressing neurons from the same neuronal culture (Fig. S3). We therefore conclude that expression of eOPN3 is well-tolerated in mammalian neurons and does not result in significant light-independent physiological changes in neuronal excitability.

Next, we tested eOPN3 in pyramidal neurons of organotypic hippocampal slice cultures, a preparation that preserves the anatomical and functional connectivity between neurons in the CA3 and CA1 regions. We co-expressed eOPN3-mScarlet with cytoplasmic mCerulean via single-cell electroporation in a small set of CA3 pyramidal neurons. One week after electroporation, both constructs were strongly expressed without affecting CA3 cell morphology (Fig. 1I). To characterize the effects of somatodendritic eOPN3-activation on neuronal excitability, we recorded from CA3 pyramidal cells expressing eOPN3 using whole-cell patch clamp electrophysiology. Applying light directly to the somatodendritic region triggered long-lasting photocurrents reversing at −105.1 ± 0.9 mV (Fig. S4A), close to the calculated K^+^ reversal potential of −102.5 mV, indicating activation of endogenous GIRK channels. This eOPN3-dependent K^+^-conductance led to a lower input resistance (Fig. S4B), a decrease in electrically evoked action potential firing (Fig. S4C), a slight hyperpolarization of the resting membrane potential (Fig. S4D) and an increased rheobase (Fig. S4E). Next, we asked whether eOPN3 can be detected in distal axons of electroporated CA3 neurons, where activation of the G_i/o_-pathway should lead to inhibition of synaptic neurotransmitter release. To verify axonal expression, we traced the axons of mCerulean-expressing CA3 pyramidal neurons to the *stratum radiatum* in CA1 (Fig. 1I). Axons and boutons expressing mCerulean consistently showed expression of eOPN3-mScarlet, indicating that the rhodopsin is present at presynaptic terminals.

### Activation of eOPN3 leads to suppression of neurotransmitter release

Our findings demonstrated that eOPN3 can reliably couple to the G_i/o_-signaling pathway, evoke GIRK-mediated currents and traffic to distal axon terminals in hippocampal neurons. We therefore asked whether activation of eOPN3 in presynaptic terminals triggers changes in neurotransmission via G-protein activation, similar to the DREADD hM4Di (Fig. S5). To address this, we recorded from cultured autaptic hippocampal neurons, in which brief depolarizations to 0 mV trigger unclamped action potentials (APs) that evoke synaptic currents in the same neuron (Bekkers & Stevens, 1991). Light delivery to eOPN3-expressing autaptic neurons resulted in a robust and long-lasting decrease of excitatory postsynaptic currents (EPSCs; Fig. 2A) and led to an increase in the paired-pulse ratio (Fig. 2B), consistent with a decrease in release probability upon eOPN3 activation (Dobrunz, et al., 1997). Light-triggered suppression of release was also found in autaptic hippocampal interneurons and was similarly accompanied by an increase in the paired-pulse ratio of the inhibitory postsynaptic currents (Fig. 2C). To determine the light sensitivity of eOPN3, we varied the light exposure between 0.2 μW·s·mm^−2^ and 20 mW·s·mm^−2^ (Fig. 2D). The half-maximal effect size was reached at 2.90 μW·s·mm^−2^, meaning that 1 s continuous illumination at 2.9 μW·mm^−2^ was sufficient to reach half maximal inhibition of synaptic vesicle release. In order to estimate the time course of the eOPN3-mediated effect on synaptic release, we elicited trains of APs at 10 Hz and applied light after 200 such APs, when EPSC amplitudes reached a steady state. We found that eOPN3-mediated suppression of release was rapid, with an onset time constant (τ_on_) of 0.24 s, and saturated after 1 s (Fig. 2E). Furthermore, activation of eOPN3 significantly decreased the frequency of AP-independent miniature EPSCs (Fig. 2F), but not their amplitude (Fig. 2G). Together, these results are consistent with a presynaptic action of this photoreceptor on neurotransmission.

**Figure 2.**
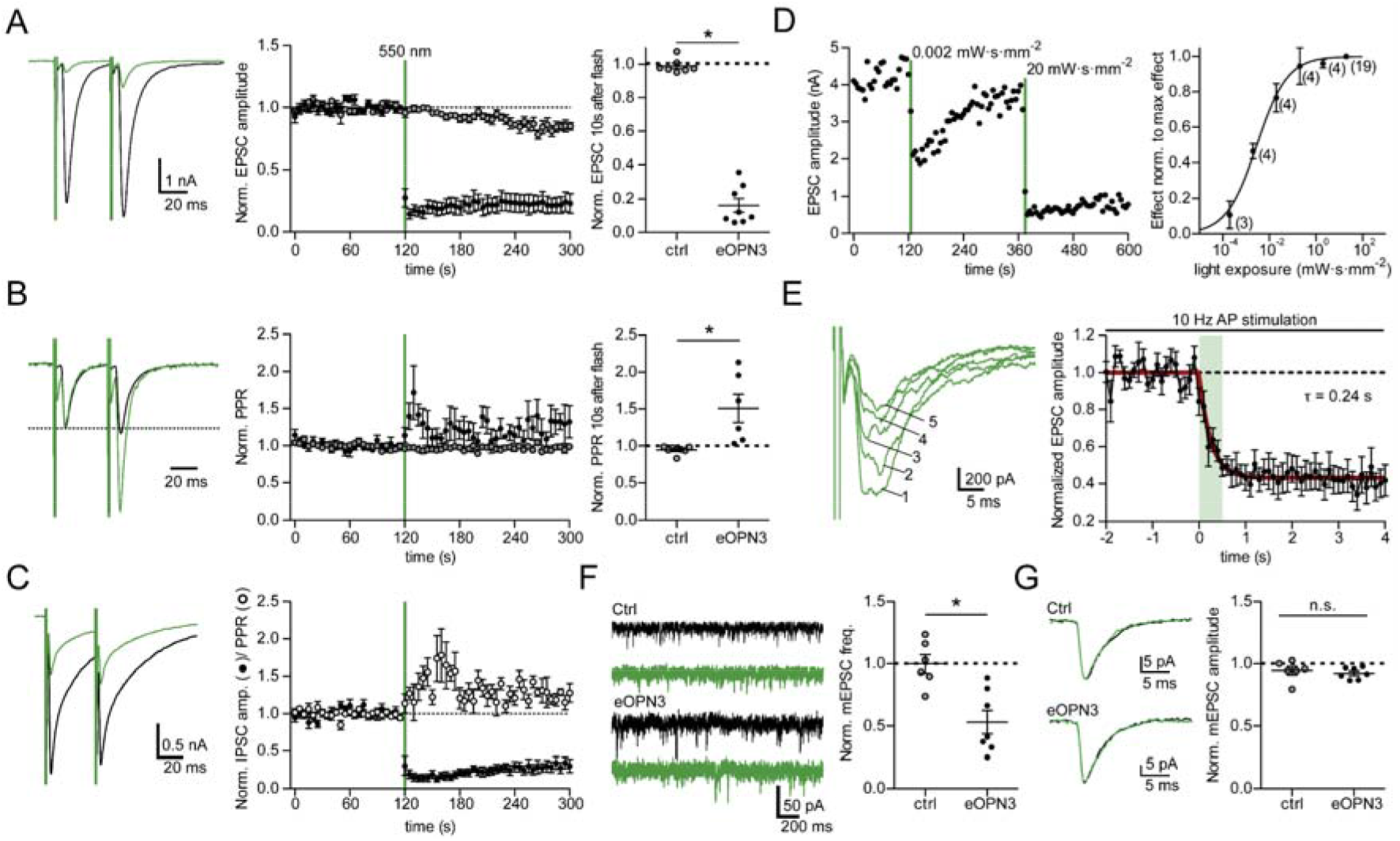
Light-induced inhibition of neurotransmitter release in autaptic hippocampal neurons expressing eOPN3. **(A)** Typical autaptic EPSCs evoked by a pair of 1 ms depolarizing current injections (40 ms inter-stimulus interval, injected currents clipped for presentation) before (black) and after (green) illumination with 550 nm light (40 mW·mm^−2^, unless otherwise indicated). Traces are averages of 6 sweeps. A 500 ms light pulse caused sustained suppression of EPSCs in eOPN3-expressing neurons. EPSCs decreased to 16 ± 4% of baseline (n = 8), while EPSCs in control neurons were not affected by illumination (open circles, n = 7, p = 3·10^−4^ two-tailed Mann-Whitney test). (**B**) Traces from (A) scaled to the amplitude of the first EPSC (dashed line). Illumination increased the paired-pulse ratio (EPSC_2_/EPSC_1_) in the eOPN3-positive neurons (n = 6) compared to controls (p = 1.2·10^−3^ unpaired, two-tailed Student’s t-test). (**C**) Similar to the effect in glutamatergic neurons, eOPN3 activation strongly suppressed IPSCs in GABAergic neurons and increased the PPR compared to the pre-light baseline (IPSCs: n = 7; PPR: n = 5). **(D)** Quantification of light exposure required for half maximal synaptic inhibition. Normalized effect size was fit as a sigmoidal dose-response curve (n is reported next to the measurement points, EC50 = 2.895 μW·s·mm^−2^). (**E**) Time-course of the eOPN3 activation on transmitter release. Light was applied after obtaining stable baseline EPSC amplitudes evoked by APs triggered at 10 Hz. Traces show five consecutive EPSCs of the train following the onset of a single 500 ms light pulse. EPSCs decreased with a time constant τ_on_ of 240 ms (n = 6). (**F**) Representative traces of mEPSCs (*left*) and quantification (*right*). eOPN3 activation decreased mEPSC frequency to 53 ± 9% compared to baseline (n = 7), significantly different from controls (n = 6, p = 3·10^−3^, two-tailed Mann-Whitney test). (**G**) Quantal amplitude was not significantly different between eOPN3-expressing and control neurons after illumination (p = 0.3 unpaired, two-tailed Student’s t-test). Plots show individual data points and average (black) ± SEM.

If the inhibitory effect of eOPN3 on synaptic transmission was mediated primarily by the G_i/o_-pathway, we predicted that light delivery and GABA_B_ receptor activation would lead to a similar reduction in EPSCs in eOPN3-expressing neurons. Moreover, pharmacological blockade of the G_i/o_ signaling pathway should reduce both GABA_B_ receptor- and eOPN3-mediated inhibition of synaptic transmission to a similar extent, as shown for hM4Di (Fig. S5C). Indeed, the effect of eOPN3 activation on synaptic transmission was similar to the effect of the GABA_B_ agonist baclofen, a potent modulator of neurotransmitter release (Fig. 3A, B; (Rost, et al., 2011; Scanziani, et al., 1992)). Pre-incubating the neurons with the G_αi/o_ subunit blocker pertussis toxin (PTX) blocked both the eOPN3 and the baclofen-mediated effects (Fig. 3A, B), indicating that eOPN3 reduced the vesicle release probability through the PTX-sensitive G_i/o_ protein signaling cascade. To examine whether the effects on synaptic transmission are dependent on GIRK channel activation, we applied SCH23390, which blocks GIRK channel currents (Kuzhikandathil & Oxford, 2002). Bath application of SCH23390 abolished the outward currents evoked by green light at the somatic compartment (Fig. 3C), but had no detectable impact on the light-activated suppression of synaptic release in the same neurons (Fig. 3D). These results suggest that the synaptic effects of eOPN3 are not mediated by blocking the propagation of APs, but rather by direct G protein-mediated effects at the presynaptic compartment (Wu & Saggau, 1994; Zurawski, et al., 2019).

**Figure 3.**
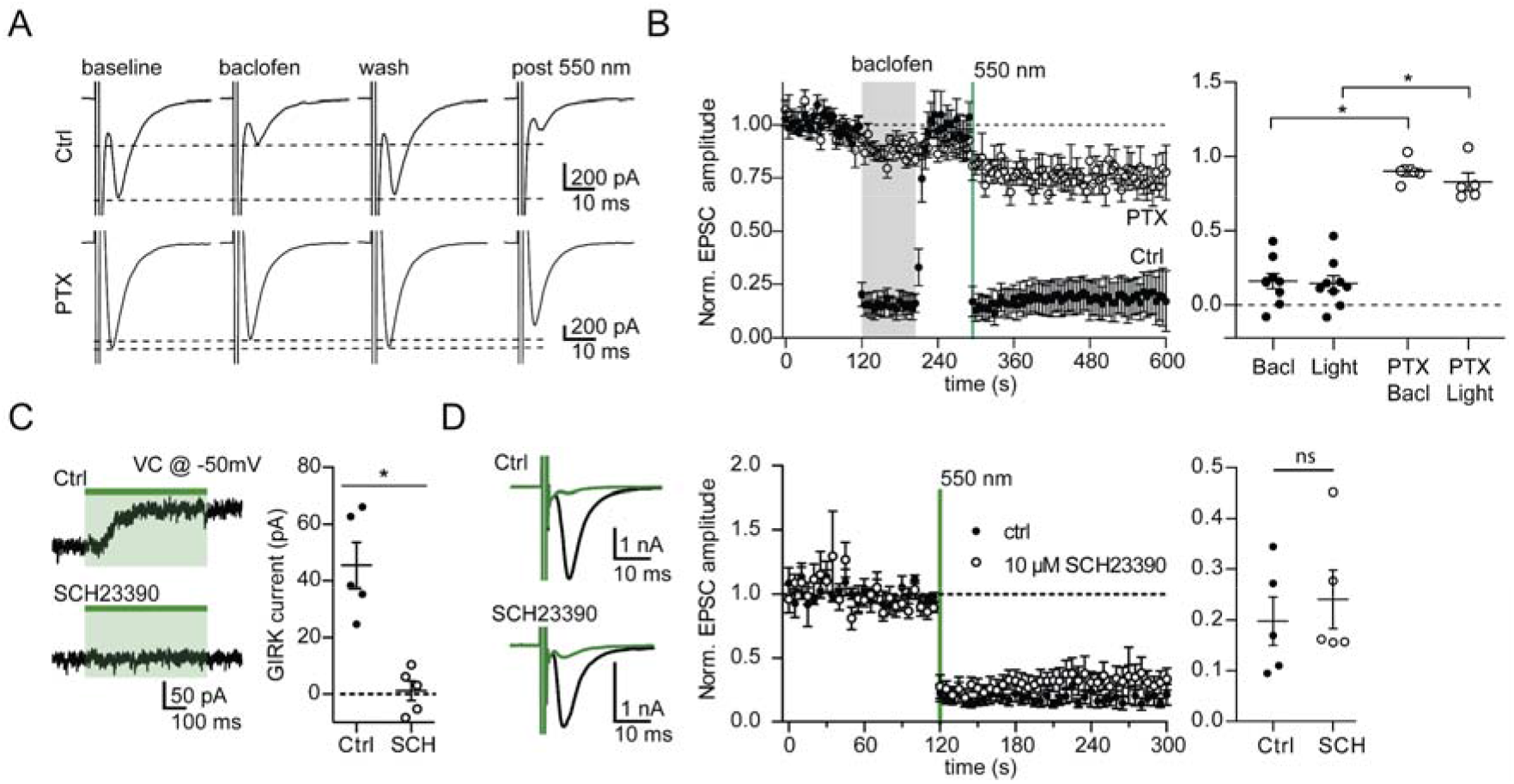
The effect of eOPN3 on neurotransmitter release is sensitive to pharmacological inhibition of G_i/o_-protein signaling but is not affected by a GIRK channel blocker. (**A**) Action potential-evoked EPSCs in control neurons (upper row) were suppressed both by the GABA_B_R agonist baclofen (30 μM) and by subsequent activation of eOPN3 with 550 nm light (500 ms, 40 mW·mm^−2^). In pertussis toxin (PTX)-treated neurons (20-26 h pre-treatment, 0.5 μg ml^−1^, bottom row), both baclofen and eOPN3 largely failed to suppress release. (**B**) Averaged time-course of EPSCs recorded in neurons treated with PTX (filled circles; n = 5) and neurons not treated with PTX (empty circles; n = 9; p = 3·10^−4^ Kruskal-Wallis test followed by Dunn’s multiple comparison tests: p<0.05 for Bacl vs PTX Bacl, Light vs PTX Bacl and Light vs PTX Light). (**C**) Illumination of eOPN3-expressing neurons evokes robust outward currents (45.5 ± 8.1 pA, n = 5), which are abolished in the presence of the GIRK channel blocker SCH23390 (10 μM, 1.2 ± 3.5 pA; n = 5; p = 1·10^−3^ unpaired, two-tailed Student’s t-test). (**D**) The extent and time-course of EPSC suppression by eOPN3 activation is not affected by the GIRK channel blocker SCH23390 (closed circles: ctrl recordings, n = 5; open circles: SCH23390, n = 5; p = 0.59 unpaired, two-tailed Student’s t-test). Plots show individual data points and average (black) ± SEM.

We next tested whether presynaptically expressed eOPN3 can be used to inhibit synaptic transmission in organotypic slices, where axon terminals can be locally illuminated independently of the neuronal soma. As described above, we expressed eOPN3 in individual hippocampal CA3 neurons using single-cell electroporation. We then recorded from pairs of CA3 and CA1 neurons, selecting only CA1 neurons that displayed evoked postsynaptic responses to paired-pulse stimulation of the recorded presynaptic CA3 cell (Fig. 4A). These recordings were performed at near-physiological temperature (33 ± 1°C) to better estimate eOPN3 kinetics under physiological conditions. Consistent with autaptic recordings, brief (500 ms), local illumination of the axonal terminals in CA1 induced a potent and long-lasting but reversible reduction of the evoked EPSC amplitude to 19 ± 4% of baseline in CA1 neurons (Fig. 4B-E and S6C-F). Light application in CA1 neither induced AP failure nor GIRK-mediated hyperpolarization in the recorded presynaptic neurons (Fig. S6A-B), suggesting that activation of eOPN3 in the axonal compartment does not reduce somatic excitability. In accordance with a reduction in evoked release and thus a direct effect of eOPN3 on neurotransmitter release from the presynaptic terminals, we found that both the coefficient of variation (CV, Fig. 4F) and the paired-pulse ratio (PPR, Fig. 4G) increased following illumination in almost all the recorded pairs. Recovery of postsynaptic currents was measured by extending the recording duration to several minutes after light illumination. The time until 50 % EPSC recovery was 6.58 ± 1.78 min (Fig. S6C-F). Synaptic transmission in non-expressing CA3-CA1 control pairs was unaffected by light stimulation (Fig. 4E-G). We therefore conclude that eOPN3 robustly activates the G_i/o_ pathway in neurons, leading to efficient suppression of presynaptic vesicle release that recovers spontaneously within minutes.

**Figure 4.**
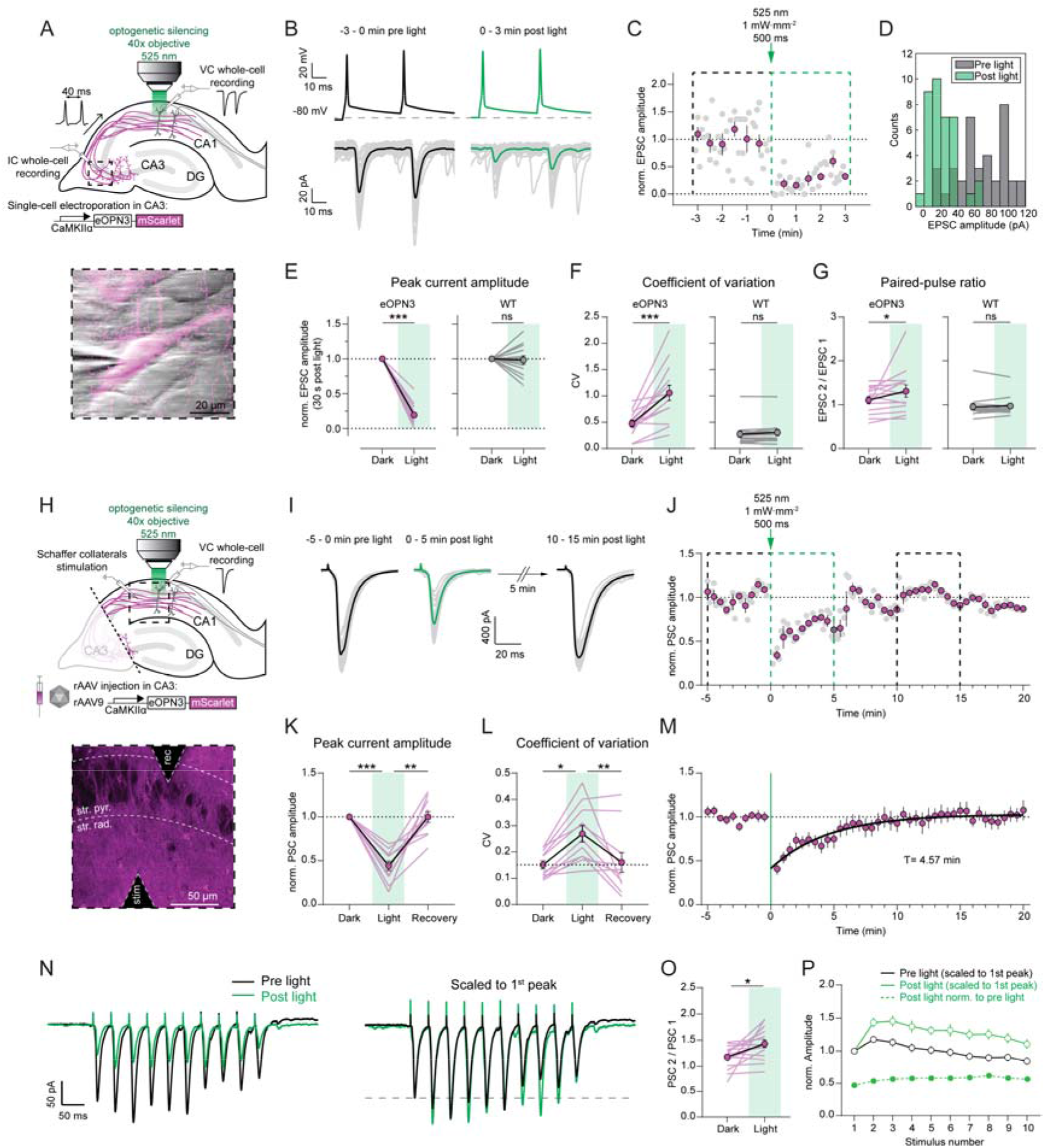
eOPN3 activation induces long-lasting, reversible inhibition of synaptic transmission at Schaffer collateral synapses. **(A)** Schematic diagram of experimental setup for whole-cell paired-recordings in organotypic hippocampal slices: pairs of APs (40 ms ISI, 0.2 Hz) were triggered in an eOPN3-transfected CA3 pyramidal neuron while recording EPSCs from a postsynaptically connected non-expressing CA1 neuron. A brief light pulse (500 ms, 525 nm, 1 mW·mm^−2^) through the objective (illuminated area = 0.32 mm^2^) in CA1 was used to activate eOPN3 locally at axon terminals innervating the postsynaptic CA1 pyramidal cell. Insert shows an IR-scanning gradient contrast image (IR-SGC) synchronized with the fluorescence images of patch-clamped, eOPN3 expressing CA3 neuron. **(B)** *Top*: representative voltage traces of electrically induced APs from an eOPN3 expressing CA3 neuron, before and after light delivery to the CA1 region (dotted line shows the resting membrane potential. Note that APs were still reliably evoked after light stimulation). *Bottom*: corresponding current traces from a postsynaptic CA1 neuron in response to the paired-pulse stimulation, before and after light delivery (gray: single trials, black and green: averaged trials). **(C)** Time course of the normalized EPSCs peak amplitudes from the example shown in B (gray dots = single trials, magenta = 30 s time bins). **(D)** Histogram count of peak current amplitudes of the example shown in B. **(E)** Quantification of presynaptic eOPN3 activation on EPSC amplitudes in the eOPN3 group (*left*) and wild-type (WT) control group (*right*) (eOPN3: 0.19 ± 0.04, n = 14 pairs from 14 slices, p = 0.0001, Wilcoxon test; WT: 0. 98 ± 0.06, n = 13 pairs from 13 slices, p = 0.4973, Wilcoxon test). **(F)** Coefficient of variation of EPSCs in the dark and after light application for the eOPN3 (*left*) and control group (*right*) (eOPN3 dark: 0.48 ± 0.06, eOPN3 light: 1.06 ± 0.15, n = 14 pairs from 14 slices, p = 0.0004, Paired t-test; WT dark: 0.27 ± 0.06, WT light: 0.31 ± 0.06, n = 13 pairs from 13 slices, p = 0.1099, Wilcoxon test). **(G)** Paired-pulse ratio change in the dark compared to after light application for the eOPN3 (*left*) and control group (*right*) (eOPN3 dark: 1.11 ± 0.08, eOPN3 light: 1.32 ± 0.14, n = 14 pairs from 14 slices, p = 0.0245, Wilcoxon test; WT dark: 0.95 ± 0.07, WT light: 0.97 ± 0.06, n = 13 pairs from 13 slices, p = 0.5879, Wilcoxon test). **(H)** Schematic diagram of experimental setup for field stimulation. Before each experiment, CA3 somata were cut off to avoid antidromic spikes and to exclude somatic effects of eOPN3 activation. Isolated CA3 axons were stimulated with a glass monopolar electrode (at 0.1 Hz) to elicit postsynaptic currents (PSCs) recorded from a non-expressing CA1 pyramidal neuron. Light was applied through the objective as described in A to activate eOPN3 in Schaffer collateral axons. Insert shows a 2-photon image (single plane) of the CA1 region with the stimulating and recording electrodes positioned in the *stratum radiatum* and *stratum pyramidale*, respectively. Schaffer collateral eOPN3-expressing axons are visible in magenta, surrounding CA1 pyramidal neurons (dark shadows). **(I)** Representative voltage traces (PSCs) before, immediately and 10 min after light (gray: single trials, black and green: average trials). **(J)** Time course of the normalized PSC peak amplitudes from the example shown in I. Dotted boxes indicate the time periods shown in I. **(K)** Quantification of eOPN3 effect on postsynaptic responses (“Dark”: 5 min period before light; “Light”: maximal eOPN3 effect during first 30 s post light, 0.44 ± 0.05, p < 1·10^−4^; “Recovery”: 10-15 min period after light, 0.99 ± 0.06, p = 0.0019; n = 11 slices, Friedman test with Dunn’s multiple comparison test). **(L)** Quantification of the effect of eOPN3 activation on the coefficient of variation. In this case, for CV calculation, “Light” refers to the 5 min post light application matching the duration of the two other conditions (“Dark”: 0.15 ± 0.02; “Light”: 0.27 ± 0.03, p = 0.0167; “Recovery”: 0.16 ± 0.04, p = 0.0085, n = 11 slices, Friedman test with Dunn’s multiple comparison test). **(M)** Summary of all field stimulation experiments. Fitted mono-exponential function is shown in black. **(N)** *Left*: representative voltage traces in response to a train stimulation consisting of 10 pulses at 25 Hz. Traces are averages of 5 sweeps each. *Right*: same traces as on the left each scaled to its 1^st^ PSC peak amplitude, showing sustained facilitation. **(O)** Quantification of the PPR (PSC 2 / PSC 1 of the train), showing increased facilitation (Dark: 1.18 ± 0.05, Light: 1.43 ± 0.07, p = 0.0105, n = 16 slices, Paired t-test). **(P)** Summary of all train stimulation experiments, showing modulation of short-term plasticity after eOPN3 activation.

To predict the effects of eOPN3-mediated inhibition *in vivo*, we replaced single-cell electroporation with virus injection in CA3 of organotypic hippocampal slice cultures. Transducing a larger portion of presynaptic cells with virus injections emulates the most commonly used method for gene transfer *in vivo* (Fig. 4H-M). To avoid both recurrent polysynaptic activity of the CA3 network and contribution of somatic eOPN3 activation, CA3 somata were removed by dissecting at the boundary of CA3 to CA1 before each experiment (Fig. 4H). Isolated Schaffer collateral axons were then stimulated with an electrode placed in the *stratum radiatum* to elicit postsynaptic currents (PSCs) in non-expressing CA1 pyramidal neurons. The PSC amplitude was attenuated by 56 ± 5% following a single 500 ms light pulse to the terminal field in the CA1 (Fig. 4I-L), and recovered to baseline levels with a time constant of 4.57 min (95% CI: 4.19 to 4.97; R^2^: 0.90). As in our paired recordings, the CV of synaptic responses increased within the 5 min following light stimulation, and eventually returned to baseline values. The lower efficacy of PSC amplitude reduction recorded in this experimental setup (Fig. 4K) compared with the efficacy observed in paired recordings (81 ± 4%, Fig. 4E) is likely due to the contribution of non-expressing axons to the PSCs evoked by field stimulation.

GPCRs may act at presynaptic terminals as canonical or non-canonical modulators of synaptic transmission (Zurawski, et al., 2019). It has been reported that canonical GPCR-mediated presynaptic inhibition decreases neurotransmission by altering the probability of vesicle release and changing the short-term plasticity profile of modulated synapses (Chalifoux & Carter, 2011), leading in some cases to suppression of initial release but facilitation of subsequent responses. To better characterize the efficacy of eOPN3-mediated synaptic inhibition during higher firing rates, we applied trains of 10 stimulations at 25 Hz (Fig. 4N-P). Postsynaptic responses in the dark showed facilitation for the initial pulses while displaying depression towards the end of the train. In accordance with our previous single-pulse field stimulation results, light activation of eOPN3 inhibited the first pulse by an almost identical amount (single pulse stimulation: 44 ± 5% vs. train stimulation: 47 ± 5% of initial strength). Consistent with our single-cell stimulation (paired CA3-CA1 recordings) data, eOPN3 increased the PPR of the initial two pulses (PSC 2 / PSC 1) and maintained facilitation throughout the train. Nonetheless, light activation of eOPN3 robustly suppressed the entire sequence of PSCs in the stimulus train albeit to a slightly lower degree for all the consecutive pulses relative to the initial one (suppression of the 10^th^ pulse was 43 ± 2 % of the initial strength).

### Integration of eOPN3-based manipulation with two-photon Ca^2+^ imaging

Experiments investigating neuronal circuits increasingly rely on two-photon imaging of optical indicators. To assess whether eOPN3 can be combined with two-photon imaging, we tested eOPN3 activation by two-photon absorption. In CA3 pyramidal cells of organotypic hippocampal cultures expressing eOPN3 and GIRK2-1, we compared green-light evoked GIRK channel currents to fast spiral scanning on the soma or slow raster scanning across the somatodendritic compartment with a femtosecond-pulsed infrared laser at wavelengths ranging from 800 to 1070 nm and at intensities ranging from 10 to 100 mW (Fig. 5A-C). Spiral scans did not evoke any detectable photocurrents (Fig. 5B). Only slow raster scans at wavelengths above 980 nm and intensities above 30 mW resulted in very small photocurrents of less than 10 pA on average (Fig. 5C). In contrast, green-light activation of eOPN3 in the same cells evoked more than 20-fold larger photocurrents (Fig. 5B). Thus, eOPN3 can be combined with two-photon imaging of blue-shifted sensors with minimal cross-activation. Previous studies have shown that different neurotransmitters and neuromodulators can act via metabotropic signaling to alter Ca^2+^ influx through Ca_v_2 voltage-gated Ca^2+^ channels located both at the presynaptic and postsynaptic compartment with different functional implications (Wu & Saggau, 1994; Ikeda, 1996; Herlitze, et al., 1996; Chalifoux & Carter, 2011; Burke, et al., 2018). At presynaptic terminals, G_i_-coupled GPCRs can suppress neurotransmitter release via G_βγ_-mediated inhibition of voltage-gated Ca^2+^ channels (Herlitze, et al., 1996; Kajikawa, et al., 2001), possibly by delaying the time of first opening or by shifting the voltage-dependency of channel activation (Bean, 1989). We therefore took advantage of the red-shifted two-photon cross section of eOPN3 and tested whether eOPN3 activation in presynaptic terminals reduces AP-evoked Ca^2+^ influx. We evoked single APs in CA3 cells co-expressing eOPN3 and jGCaMP7f (Dana, et al., 2019), while imaging the corresponding presynaptic Ca^2+^ transients in CA3 cell axonal boutons in CA1 *stratum radiatum* (Fig. 5D,E). The GIRK channel blocker SCH23390 was added to exclude potentially confounding GIRK channel-mediated hyperpolarization effects. Green light pulses locally applied to the CA1 region before each trial significantly reduced presynaptic Ca^2+^ influx in a GIRK-independent manner (Fig. 5F-G), indicating that eOPN3 acts directly at voltage-dependent Ca^2+^ channels at presynaptic terminals similar to native G_i_-coupled receptors.

**Figure 5.**
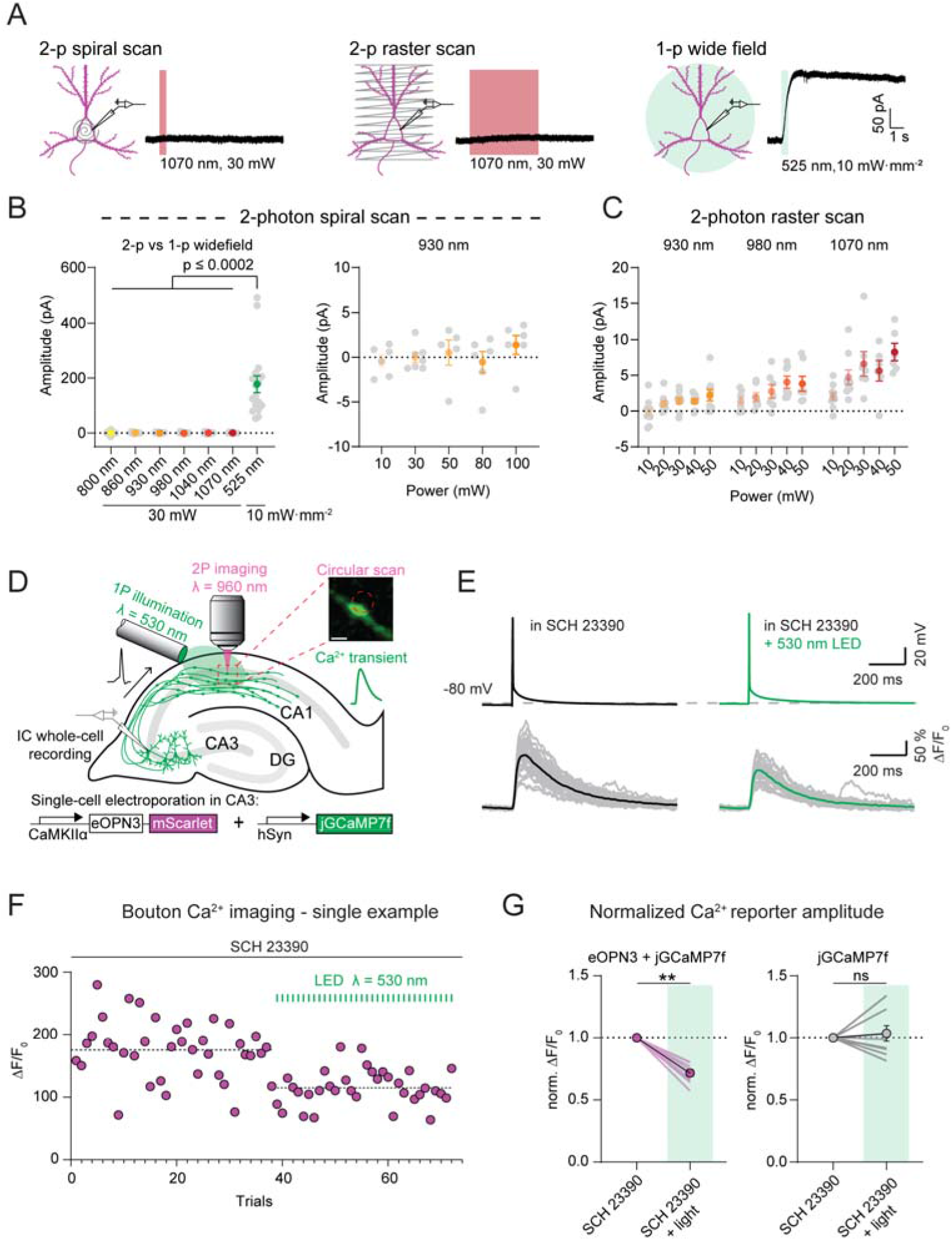
eOPN3 2-photon activation properties and modulation of presynaptic voltage-gated Ca^2+^ channels. **(A)** Two-photon versus single-photon activation of eOPN3 in CA3 pyramidal neurons in organotypic hippocampal slice cultures expressing eOPN3-mScarlet and GIRK2-1. Somatic 500 Hz spiral scans (2 ms/spiral, 250 cycles, 500 ms total duration) or raster scans (FOV = 106*106 μm, 512×512 pixels, 1.8 ms/line, 5 frames, 4.6 s total duration) at 1.09 Hz over the somatodendritic compartment were used for two-photon activation characterization. Wide-field stimulation with green light was used for subsequent single-photon activation of eOPN3 in the same neuron. Example voltage-clamp traces show photocurrents obtained by the different stimulation modalities in the same cell. **(B)** Quantification of the photocurrents elicited by two-photon versus single-photon illumination. *Left*: excitation wavelength used for spiral scanning of eOPN3 ranged from 800 nm to 1070 nm, each at 30 mW. None of the stimulation parameters resulted in notable photocurrents. Illumination with green light, on the other hand, produced significant photocurrents (Kruskal-Wallis test, Dunn’s multiple comparisons test). *Right*: Increasing laser intensity during spiral scans at 930 nm did not result in significant photocurrent. Typical Ca^2+^ imaging experiments at 930 nm require laser intensities <30 mW. **(C)** Slower and longer raster scanning over a larger field of view resulted in minimal outward currents and was wavelength and laser-intensity dependent. **(D)** Schematic diagram of presynaptic Ca^2+^ imaging experiments. Single action potentials were triggered via a patch pipette in a CA3 pyramidal neuron co-expressing eOPN3 and jGCaMP7f or jGCaMP7f alone as control while evoked Ca^2+^ influx at distal presynaptic terminals in *stratum radiatum* of CA1 was monitored by two-photon microscopy. Insert shows a single-plane jGCaMP7f image of an en passant bouton and the circular imaging-laser scanning path (red dashed circle, scale bar = 1 μm). A fiber-coupled LED (λ = 530 nm; 83 μW at the fiber tip) was used to locally activate eOPN3 in CA1 the presence of the GIRK channel blocker SCH 23390. **(E)** Top: representative voltage traces of electrically evoked action potentials in a transfected CA3 pyramidal neuron in the dark and after a green light pulse (dashed line shows the resting membrane potential). Bottom: corresponding Ca^2+^ responses from a presynaptic bouton. Single trails are shown in gray, black and green traces represent the averaged responses before and after light, respectively. **(F)** Single bouton Ca^2+^ imaging experiment showing jGCaMP7f peak transients in the dark (gray circles) and after green light pulses (green circles) indicating a light-dependent decrease in presynaptic Ca^2+^ influx. Dotted lines show the average for the two conditions. **(G)** Quantification of normalized eOPN3-jGCaMP7f transients (*left*) (SCH 23390 + light = 0.72 ± 0.026, p = 0.002, Wilcoxon-test, n = 10 slices) and jGCaMP7f alone (*right*) (SCH 23390 + light = 1.04 ± 0.06, p = 0.8852, Paired t-test, n = 10 slices). Plots show individual data points (lines), and average (circles) ± SEM.

### *In vivo* characterization of eOPN3 mediated terminal inhibition

Next, we examined the efficacy and kinetics of eOPN3-mediated presynaptic silencing *in vivo* at the electrophysiological level. We chose to modulate the visual thalamocortical pathway, since projections from the lateral geniculate nucleus of the thalamus (LGN) are the main feed-forward input from the retina to V1 and the visual responses of V1 neurons depend on the input from LGN ( (Niell & Stryker, 2008; Froudarakis, et al., 2019). To render thalamocortical projections light controllable, we injected eOPN3-encoding virus bilaterally into the LGN (Fig. S7A). During viral expression, mice were accustomed to head fixation. 7-10 weeks after viral injection, we performed bilateral craniotomies and inserted a multi-shank silicon probe into each hemisphere’s V1 to perform extracellular recordings of local brain activity. We then probed LGN input to V1 every 30 s by presenting awake, head-fixed mice with a 4 s compound visual stimulus (Fig. S7B). Visual stimulation led to reliable evoked responses in V1 (Fig. S7C,D left). A subset of units showed an increase in their average firing rates during visual stimulus presentation (Fig. S7D). After 10 trials of visual stimulus presentation, we activated eOPN3 in LGN terminals unilaterally by 30 s continuous illumination (2 mW at the fiber tip) directed at V1. eOPN3 activation resulted in a reduced impact of visual stimulation on the V1 network activity (Fig. S7C,D right), with responsive units reducing the response amplitude (Fig. S7E). To allow for a robust characterization of eOPN3 mediated vesicle release inhibition recovery, we selected units that showed visually-evoked responses that were suppressed by more than 50% during eOPN3 activation (14 of 54 units). The average response amplitude of these units recovered with a time constant of 5.17 min (95% CI: 1.12 to 7.20 min; R^2^: 0.82), showing that eOPN3 can be utilized to inhibit synaptic vesicle release *in vivo* robustly and reversibly (Fig. S7F). Note that units recorded simultaneously at the contralateral (non-illuminated) side did not show a change in their visual stimulus presentation response after eOPN3 activation on the ipsilateral hemisphere, demonstrating the spatial specificity of the manipulation.

To examine the efficacy and kinetics of eOPN3-mediated presynaptic silencing *in vivo* on the behavioral level, we used eOPN3 to inhibit dopaminergic (DA) input to the dorsomedial striatum (DMS) of mice during free locomotion. Previous work has demonstrated the important role of nigrostriatal DA projections in the control of animal locomotion (Alcaro, et al., 2007; Kravitz, et al., 2010; Grealish, et al., 2010; Tecuapetla, et al., 2014; Barter, et al., 2015; Borgkvist, et al., 2015; Silva, et al., 2018). Briefly, striatal D1-expressing medium spiny neurons (MSNs) facilitate motion upon selective, bilateral activation, and induce a contralateral rotation upon unilateral stimulation. Conversely, D2-expressing MSNs decrease motion, and upon unilateral stimulation induce ipsilateral rotation. While D1 and D2 neurons drive motion in opposite directions, their common substantia nigra pars compacta (SNc) dopaminergic input stimulate D1-expressing MSNs, while inhibiting D2-expressing MSNs. Overall, these studies suggest that unilateral inhibition of nigrostriatal DA projections would introduce an ipsiversive bias in free locomotion (Fig. 6A). We thus expressed an eOPN3- or an eYFP-expressing control vector unilaterally in dopaminergic neurons of the SNc and implanted an optical fiber above the ipsilateral DMS to allow illumination of nigrostriatal DA projections (Fig. 6B). Following at least 6 weeks of recovery, mice were placed in a square open field arena, and their behavior was tracked using DeepLabCut (Mathis, et al., 2018). Following a 10-minute baseline period, we delivered 500-ms light pulses (540 nm, 10 mW from the fiber tip) at 0.1 Hz for 10 minutes to activate eOPN3 in presynaptic DA terminals, followed by 10 minutes with no light delivery to allow for eOPN3 recovery. While roaming freely in the large arena, eOPN3-expressing mice exhibited an ipsiversive preference upon illumination (Fig. 6C, D), indicating that eOPN3 can effectively inhibit nigrostriatal projections with minimal light delivery. The rotational preference was not observed during the baseline period and became evident within the first minute following light onset. Importantly, the behavioral preference for ipsiversive turning recovered within <10 minutes of the last light pulse (Fig. 6E), in line with the recovery kinetics of eOPN3 observed in our experiments *in vitro* and *in vivo* (Figs. 4M, S6C-F and S7F). Control mice did not show such side bias or light-induced equivalent dynamics (Fig. 6C-E). Apart from their strong side preference, eOPN3 mice did not differ from control mice in distance traveled (p = 0.54, Kruskal-Wallis test), center entries (p = 0.99, Kruskal-Wallis test), or time in center (p = 0.69, Kruskal-Wallis test). The magnitude of the observed behavioral effect of eOPN3 activation, quantified as the rotation index (Fig. 6D, *insets*; see Methods), was positively correlated with expression levels across individual mice (p=6.1·10^−3^, R^2^=0.81), during the light activation period, but not before light delivery or after its termination (Fig. 6F). No significant correlation was found with the average velocity before, during or after eOPN3 activation (Fig. 6F). Finally, one week after the initial test, we repeated the test using the same parameters. We found a high correlation in the light evoked rotational bias between the first and second trial in each mouse (Pearson’s correlation coefficient: 0.8147; *p* = 0.0256). Taken together, our results demonstrate that eOPN3 can be used for synaptic terminal inhibition in behaving animals, with high light-sensitivity, precisely timed onset, and behaviorally relevant recovery time.

**Figure 6.**
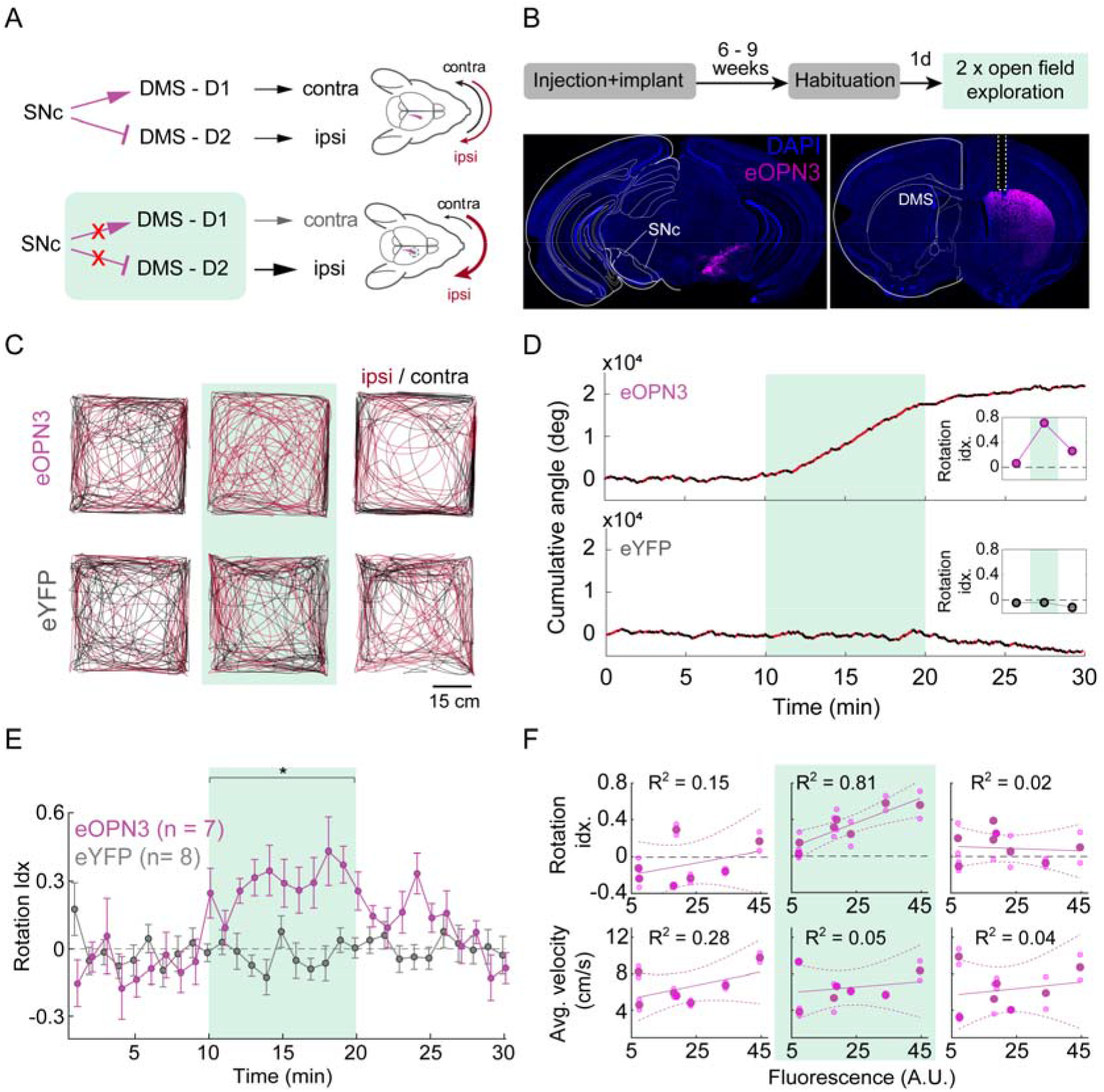
eOPN3-mediated suppression of dopaminergic projections from the substantia nigra to the dorsomedial striatum leads to ipsiversive bias during free locomotion. **(A)** Schematic diagram of the experimental setup and hypothesis. Unilateral expression of eOPN3 in SNc dopaminergic neurons and light-mediated suppression of their striatal projections would lead to an ipsiversive side bias during free locomotion **(B)** Top: experimental timeline. Bottom: Representative images of neurons expressing eOPN3-mScarlet in the SNc (*left*) and their striatal projections (*right*) in DAPI-stained brain sections. **(C)** Locomotion paths of representative eOPN3 (*top*) and eYFP (*bottom*) mice, over successive 10-minute periods: (*left* to *right*) before, during and after light delivery (540 nm light pulses, 10 mW from the fiber tip, 540 nm at 0.1 Hz), together covering continuous 30 minutes sessions. Red and black color code path segments where the mice showed ipsilateral or contralateral angle gain, respectively. **(D)** Representative cumulative angle traces of individual eOPN3-expressing (*top*) and eYFP-expressing (*bottom*) mice, over 30 minutes of free locomotion in an open field arena. Red and black colors depict ipsilateral or contralateral segments, respectively. Green shaded region marks the light delivery period. **(E)** The rotation index (mean ± SEM), calculated as the difference between cumulative ipsilateral and contralateral rotation, divided by their sum, over 1-minute bins for eOPN3-expressing mice (magenta, *n* = 7) and eYFP controls (gray, *n* = 8). Green shaded region marks the light delivery period, where eOPN3 demonstrate significant ipsiversive bias (p = 1.3·10^−3^ Kruskal-Wallis test followed by Bonferroni-Holm corrected pairwise comparisons using Wilcoxon rank sum tests. Baseline: ctrl vs. eOPN3 p = 1; light: ctrl vs. eOPN3 p = 1.9·10^−3^; post light: ctrl vs. eOPN3 p = 0.09). **(F)** Top: rotation index, calculated for individual mice before (*left*), during (*middle*) and after (*right*) light-induced activation of eOPN3, plotted against eOPN3 expression levels measured at the DMS projections (symbols). Solid and dashed lines are linear regression fit with 95% confidence intervals, respectively. Bottom: average velocity of individual mice, plotted against expression levels in the same manner shown above. R^2^ values are indicated separately for each plot.

## Discussion

Optogenetic silencing is a powerful tool for functionally dissecting neuronal circuits and understanding the contribution of defined neuronal populations to behavioral processes. However, silencing of long-range axonal projections has posed a formidable challenge. This is due to an inefficacy of most optogenetic tools to suppress synaptic transmission and to paradoxical effects of others (Mahn, et al., 2016). Suppression of axonal APs with potassium-conducting optogenetic tools (Cosentino, et al., 2015; Alberio, et al., 2018; Bernal Sierra, et al., 2018; Beck, et al., 2018) has not been shown to be effective for presynaptic vesicle release inhibition. Chemogenetic tools such as hM4Di can be used for silencing presynaptic release (Stachniak, et al., 2014), but suffer from slow kinetics due to the unbinding and clearance of their small-molecule ligands. Some newly developed optogenetic tools have also been used to selectively suppress exocytosis (Liu, et al., 2019), but these tools necessitate protein turnover to reinstate synaptic transmission and are consequently intrinsically slow. We therefore sought to harness the efficacy of G_i/o_-coupled signaling and combine it with the advantages of optogenetic tools, to engineer an effective presynaptic silencing tool of high spatiotemporal precision. Such optogenetic tools would induce similar effects as G_i/o_-coupled DREADDs but without the need for infusion of small-molecule ligands into the targeted brain regions, thereby allowing optogenetic silencing of neurotransmitter release through direct illumination of synaptic terminals.

Our results demonstrate that a mosquito homolog of encephalopsin (OPN3) can indeed selectively recruit G_i/o_ signaling in mammalian neurons. Optimization of this rhodopsin (yielding eOPN3) led to enhanced membrane targeting and improved expression in long-range axons. In autaptic neuron cultures, the inhibitory effects of eOPN3 on synaptic transmission were identical to those of the activation of endogenous GABA_B_ receptors, suggesting that a common mechanism underlies this effect. One potential caveat to the use of G_i/o_-mediated inhibition for the manipulation of neuronal and synaptic activity is that the biochemical signaling pathways and the effector proteins might differ among cell types and subcellular compartments. In our experiments, however, we observed robust inhibitory effects of eOPN3 in various types of glutamatergic, GABAergic and dopaminergic neurons. In the somatic compartment of neurons expressing eOPN3, we observed light-triggered GIRK channel-mediated currents which could be further enhanced by co-expressing GIRK2-1 channels. In hippocampal slices, the effect of eOPN3 activation on the intrinsic excitability of expressing neurons was relatively weak. This suggests that activation of eOPN3 in the somatodendritic compartment will lead to less efficient inhibition of neuronal spiking, compared to other K^+^ channel-mediated optogenetic silencing approaches (Bernal Sierra, et al., 2018; Beck, et al., 2018). If exclusive illumination of the axonal compartment is not feasible in a given pathway and a reduced excitability of the soma needs to be prevented, the somatodendritic effects of eOPN3 could likely be further diminished by the addition of a selective somatodendritic endocytosis sequence to the eOPN3 coding sequence (Fairless, et al., 2008; Stachniak, et al., 2014). By contrast, silencing of synaptic transmission with eOPN3 was highly efficient and independent of GIRK channel activity, suggesting that eOPN3-mediated synaptic inhibition occurs through direct activity on the highly-conserved presynaptic release apparatus and on Ca^2+^ channel function (Dittman & Regehr, 1996; Kajikawa, et al., 2001; Sakaba & Neher, 2003; Zurawski, et al., 2019). This is consistent with our observation of GIRK channel-independent suppression of spike-evoked Ca^2+^ transients after eOPN3 activation. Thus, if locally activated at synaptic terminals, eOPN3 is a robust and broadly applicable optogenetic tool for inhibition of synaptic neurotransmission, similar to the DREADD receptor hM4Di, which has been successfully used for presynaptic silencing in a variety of neuronal cell types and systems (Stachniak, et al., 2014; Evans, et al., 2018; Malvaez, et al., 2019).

The effects of GPCRs on presynaptic neurotransmitter release have been partially attributed to G-protein modulation of presynaptic Ca^2+^ influx (Herlitze, et al., 1996). Meanwhile, non-canonical presynaptic GPCR modulators have been shown to decrease the vesicle release probability without a concomitant change in short term plasticity, through Ca^2+^-dependent and independent mechanisms (Hamid, et al., 2013; Burke, et al., 2018). Our paired-pulse facilitation results, both from autaptic and organotypic cultures, suggest that eOPN3 act as a canonical presynaptic GPCR modulator: while it consistently suppresses synaptic transmission evoked during trains of action potentials, it inhibits the first pulse more strongly than it does the consecutive pulses (Fig. 4N-P). This could be due to presynaptic Ca^2+^ accumulation (Jackman & Regehr, 2017) and a depolarization-triggered relief of the G-protein interaction with voltage gated Ca^2+^ channels (Currie, 2010). Thus, eOPN3 activation biases short-term synaptic plasticity towards short-term facilitation.

We have previously shown that current approaches utilizing ion pumps for vesicle release inhibition are either not able to effectively silence presynaptic release for extended time periods or lead to alkalization of the presynaptic compartment and an increase in spontaneous neurotransmission, which can have undesired effects on the downstream target (Mahn, et al., 2016; Lafferty & Britt, 2020). Importantly, bistable rhodopsins such as eOPN3 cannot replace ion-pumping type-I rhodopsins in the range of sub second precise control over vesicle release. eOPN3 should therefore be utilized for experiments that require vesicle release inhibition in the range of minutes to hours. For even longer inhibition periods, tools such as the photoactivatable botulinum neurotoxin are likely also suitable (Liu, et al., 2019). Silencing synaptic transmission using hM4Di with local agonist infusion at the terminal field as mentioned above (Stachniak et al. 2014) should in principle allow for similar efficiency compared to eOPN3. However, eOPN3 has the advantage of more precise temporal control and reduced problems with agonist microinfusion such as potential off-site effects due to leakage to the cerebrospinal fluid. The time course of recovery after eOPN3 activation that we observed *in vitro* (Fig. 4M, Fig. S6C-F) and *in vivo* (Fig. 6E, Fig. S7F) is consistent across the four preparations and three cell types used. However, we would like to emphasize that the exact time constants will depend on cell type and expression level and should ideally be determined experimentally in every preparation.

Our *in vitro* experiments showed that eOPN3 is highly light-sensitive (Fig. 2D), likely due to its recovery kinetics. By relaxing the limitations imposed by tissue heating *in vivo*, eOPN3 allows for optical access to large brain volumes, a major constraint of type-I rhodopsins such as NpHR and Arch (Stujenske, et al., 2015; Owen, et al., 2019). In our single-photon excitation experiments, we used light exposures above 0.5 mW·s·mm^−2^ leading to complete eOPN3 activation. This approach was aimed at achieving the maximal vesicle release inhibition, making the effect of light exposures comparable as long as they are beyond saturation while not leading to tissue heating. However, for experiments where subsets of postsynaptic targets need to be specifically inhibited, light exposure should be minimized to prevent inadvertent eOPN3 activation in neighboring areas. Furthermore, the high light sensitivity of eOPN3 necessitates working in light shielded conditions when using *in vitro* preparations or transparent organisms. For behavioral experiments, we used single light pulses spaced at 0.1 Hz. The exact irradiance and duty cycle in such experiments should be calibrated based on the volume of the targeted terminal field, and the distance from other projections and somata that should remain unaffected.

We also show that eOPN3 has a very small two-photon absorption cross section at the typical wavelength ranges used for two-photon Ca^2+^ indicator imaging (Fig. 5B). Even continuous raster scanning on the soma and proximal dendrites of neurons expressing eOPN3 and GIRK2-1 only led to a mild somatic hyperpolarization, indicating that eOPN3 is not effectively activated. A potential use case would be to image the activity of a local network before and during inhibition of a given afferent via eOPN3 activation. Here, one potential concern is that the slow recovery kinetics of eOPN3 might lead to an accumulation of G_i/o_ signaling over time, even with the low two-photon absorption properties of eOPN3. This certainly warrants careful controls, but we do not expect this to represent a major constraint in classical raster scanning two-photon imaging. Typical experiments in which network activity is continuously imaged involve a much larger field of view (1×1 mm vs. 106×106 μm used here). This effectively reduces the irradiance per illuminated presynaptic terminal. Secondly, whatever activation of eOPN3 molecules does take place, it will be limited to the imaging plane, meaning that out-of-focus eOPN3 molecules will not be affected. In contrast, combination of eOPN3-mediated inhibition with scanless two-photon approaches, such as temporal focusing or holographic imaging, might lead to an increased crosstalk. Although we did not observe such an effect in our experiments, one should also take into account that eOPN3 can potentially be activated by the emission light of the imaged indicator. In both types of experiments, the imaging parameters should be optimized to minimize such cross-activation.

To the best of our knowledge this study, along with the adjoining manuscript from the Bruchas and Gereau labs using the lamprey parapinopsin (PPO; Copits et al., bioRχiv 2021) are the first to describe an optogenetic application of bistable nonvisual rhodopsins for efficient light-gated silencing of synaptic transmission. The unique spectral features of eOPN3 and PPO, particularly in their two-photon cross sections, will potentially allow them to be utilized in concert for dual-channel optogenetic control of intracellular signaling. These two rhodopsins are part of a widespread family of non-visual rhodopsins, some of which have been shown to similarly couple to G_i/o_ signaling when expressed heterologously (Koyanagi & Terakita, 2014). Thus, additional members of this rhodopsin family could potentially serve as effective tools for controlling the activity of presynaptic terminals and might be further engineered for spectral tuning or G-protein coupling specificity. Further work is needed to examine the functional properties of these little-explored photoreceptors and adapt them for optogenetic applications. Nevertheless, eOPN3-mediated silencing of transmitter release constitutes a much-needed experimental approach for light-triggered suppression of neuronal communication in the target area of long-range projections, and we expect its application will facilitate research in a variety of neurobiological studies.

## Methods

KEY RESOURCES TABLE.

KEY RESOURCES TABLE
| REAGENT or RESOURCE | SOURCE | IDENTIFIER |
| --- | --- | --- |
| Bacterial and Virus Strains |  |  |
| rAAV2/1&2.CamKIIα(0.4).OPN3-mScarlet | This Paper | N/A |
| rAAV2/1&2.CamKIIα.eYFP.WPRE | This Paper | N/A |
| rAAV2/1&2.CamKIIα(0.4).eOPN3-mScarlet | This Paper | https://www.addgene.org/Ofer_Yizhar/ |
| rAAV2/1&2.hSyn.SIO-eOPN3-mScarlet | This Paper | https://www.addgene.org/Ofer_Yizhar/ |
| Chemicals, Peptides, and Recombinant Proteins |  |  |
| (R)-baclofen | Tocris | Cat#0796 |
| Clozapine-N-Oxide | Enzo Life Science | Cat#-BML-NS105 |
| CPPene | Tocris | Cat#1265 |
| Gabazine | Tocris | Cat#1262 |
| NBQX | Tocris | Cat#1044 |
| Pertussis toxin | Sigma-Aldrich | Cat#516560 |
| Picrotoxin | Tocris | Cat#1128 |
| SCH23390 | Tocris | Cat#0925 |
| Critical Commercial Assays |  |  |
| GloSens cAMP Assay | Promega | Cat#E1171 |
| Experimental Models: Cell Lines |  |  |
| HEK293T | Sigma-Aldrich | Cat#12022001 |
| Experimental Models: Organisms/Strains |  |  |
| Mouse: C57BL/6J | Envigo | Cat#057 |
| Mouse: C57BL/6N | Envigo | Cat#044 |
| Rattus norvegicus: Sprague-Dawley | Envigo | Cat#002 |
| Rattus norvegicus: Wistar | Charles River, bred in the animal facility, UKE Hamburg | Cat#003 |
| Recombinant DNA |  |  |
| pAAV-CaMKIIα(0.4)-OPN3-mScarlet | This Paper | N/A |
| pAAV-CaMKIIα(0.4)-PufTMT-mScarlet | This Paper | N/A |
| pAAV-CaMKIIα(0.4)-OPN3-M4-mScarlet | This Paper | N/A |
| pAAV-CaMKIIα(0.4)-PufTMT-M4-mScarlet | This Paper | N/A |
| pAAV-CaMKIIα-eYFP | Karl Deisseroth, unpublished | Addgene: Cat#105622 |
| pcDNA3.1-GIRK2-1 | Eitan Reuveny | NM_001025584.2 |
| pcDNA3.1-mCerulean | Dave Piston; Rizzo et al., 2004 | Addgene: Cat#15214 |
| pAAV-CaMKIIα(0.4)-eOPN3-mScarlet | This Paper | https://www.addgene.org/Ofer_Yizhar/ |
| pAAV-hSyn-SIO-eOPN3-mScarlet | This Paper | https://www.addgene.org/Ofer_Yizhar/ |
| Software and Algorithms |  |  |
| Fiji | Schindelin, et al., 2012 | N/A |
| Matlab 2018b | Mathworks | N/A |
| RStudio Desktop | RStudio | N/A |
| Ephus | Suter, et al., 2010 | N/A |
| ScanImage | Vidrio Technologies | 2017b |
| EthoVision XT 11.5 | Noldus | N/A |

### Animals

Animal experiments were carried out according to the guidelines stated in directive 2010/63/EU of the European Parliament on the protection of animals used for scientific purposes. Animal experiments at the Weizmann Institute were approved by the Weizmann Institute Institutional Animal Care and Use Committee (IACUC); experiments in Berlin were approved by local authorities in Berlin and the animal welfare committee of the Charité – Universitätsmedizin Berlin, Germany. Experiments in Hamburg were done in accordance with the guidelines of local authorities and Directive 2010/63/EU.

### Molecular cloning of bistable rhodopsin constructs

The genes encoding mScarlet (Bindels, et al., 2016), OPN3, PufTMT, OPN3-M4 and PufTMT-M4 were synthesized using the Twist gene synthesis service (Twist Bioscience, USA). The Rho1D4 sequence (TETSQVAPA) was added at the C-terminus of all rhodopsins. All genes were subcloned into pAAV vectors under the CamKIIα promoter and in-frame with mScarlet at the C-terminus. The eOPN3 plasmid was generated by adding the Kir2.1 membrane trafficking signal (KSRITSEGEYIPLDQIDINV) between the OPN3 and the mScarlet coding sequences and the Kir2.1 ER export signal (FCYENEV) following the C-terminus of mScarlet. eOPN3 constructs and viruses are available from Addgene: https://www.addgene.org/Ofer_Yizhar/

### Production of recombinant AAV vectors

HEK293 cells were seeded at 25%-35% confluence. The cells were transfected 24 h later with plasmids encoding AAV rep, cap of AAV1 and AAV2 and a vector plasmid for the rAAV cassette expressing the relevant DNA using the PEI method (Grimm, et al., 2003). Cells and medium were harvested 72 h after transfection, pelleted by centrifugation (300 g), resuspended in lysis solution ([mM]: 150 NaCl, 50 Tris-HCl; pH 8.5 with NaOH) and lysed by three freeze-thaw cycles. The crude lysate was treated with 250 U benzonase (Sigma) per 1 ml of lysate at 37°C for 1.5 h to degrade genomic and unpackaged AAV DNA before centrifugation at 3,000 g for 15 min to pellet cell debris. The virus particles in the supernatant (crude virus) were purified using heparin-agarose columns, eluted with soluble heparin, washed with phosphate buffered saline (PBS) and concentrated by Amicon columns. Viral suspension was aliquoted and stored at –80°C. Viral titers were measured using real-time PCR. In experiments that compared between different constructs, viral titers were matched by dilution to the lowest concentration. AAV vectors used for neuronal culture transduction were added 4 days after cell seeding. Recordings were carried out between 4-20 days after viral transduction. The following viral vectors were used in this study:

AAV2/1&2.CamKIIα(0.4).OPN3-mScarlet, AAV2/1&2.CamKIIα(0.4).eOPN3-mScarlet,
AAV2/5.CamKIIα(0.4).eOPN3-mScarlet, AAV2/9.CamKIIα(0.4). eOPN3-mScarlet
AAV2/1&2.CamKIIα.eYFP.WPRE, AAV2/1&2.hSyn.SIO-eOPN3-mScarletAAV2/1&2.EF1a.DIO.eYFP.WPRE.

### Primary hippocampal neuron culture

Primary cultured hippocampal neurons were prepared from male and female P0 Sprague-Dawley rat pups (Envigo). CA1 and CA3 were isolated, digested with 0.4 mg ml^−1^ papain (Worthington), and plated into a 24-well plate at a density of 65,000 cells per well, onto glass coverslips pre-coated with 1:30 Matrigel (Corning). Cultured neurons were maintained in a 5% CO_2_ humidified incubator in Neurobasal-A medium (Invitrogen) containing 1.25% fetal bovine serum (FBS, Biological Industries), 4% B-27 supplement (Gibco), and 2 mM Glutamax (Gibco). To inhibit glial overgrowth, 200 μM fluorodeoxyuridine (Sigma) was added after 4 days of *in vitro* culture (DIV).

Neurons were transfected using the Ca^2+^ phosphate method (Graham & Eb, 1973). Briefly, the medium of primary hippocampal neurons cultured in a 24 well plate was collected and replaced with 400 μl serum-free modified eagle medium (MEM, ThermoFisher scientific). 30 μl transfection mix (2 μg plasmid DNA and 250 μM CaCl_2_ in HBS at pH 7.05) were added per well. After 1 h incubation the cells were washed 2 times with MEM and the medium was changed back to the collected original medium. Cultured neurons were used between 14 – 17 DIV for experiments. The following plasmids were used in this study: pAAV-CamKIIα(0.4)-OPN3-mScarlet, pAAV-CamKIIα(0.4)-eOPN3-mScarlet, pAAV-CamKIIα(0.4)-PufTMT-mScarlet, pAAV-CamKIIα(0.4)-OPN3-M4-mScarlet, pAAV-CamKIIα-(0.4)PufTMT-M4-mScarlet, pAAV-CamKIIα(0.4)-eYFP. The pcDNA3.1-GIRK2-1 plasmid was a gift from Eitan Reuveny.

Autaptic cultures of primary hippocampal neurons on glia cell micro-islands were prepared from newborn C57/BL6-N mice of either sex as previously described (Rost, et al., 2015). Briefly, 300 mm diameter spots of growth permissive substrate consisting of 0.7 mg ml^−1^ collagen and 0.1 mg ml^−1^ poly-D-lysine was applied with a custom-made stamp on coverslips coated with a thin film of agarose. Astrocytes were seeded onto the glass coverslips and were allowed to proliferate in Dulbecco’s modified eagle medium (DMEM) supplemented with 10% fetal calf serum and 0.2% penicillin/streptomycin (Invitrogen) for one more week to form glia micro-islands. After changing the medium to Neurobasal-A supplemented with 2% B27 and 0.2% penicillin/streptomycin, hippocampal neurons prepared from P0 mice were added at a density of 370 cells cm^−2^. Neurons were infected with AAVs at DIV 1–3 and recorded between DIV 14 and DIV 21.

### Confocal imaging and quantification

Primary cultured hippocampal neurons were transfected at 5 DIV with plasmids encoding a rhodopsin protein (mScarlet, OPN3, PufTMT, OPN3-M4, PufTMT-M4, eOPN3) along with pAAV-CamKIIα-eYFP. Four days after transfection, cells were fixed and permeabilized, washed 4 times with PBS and stained for 3 min with DAPI (5 mg/ml solution diluted 1:30,000 prior to staining). Coverslips were then mounted using PVA-DABCO (Sigma) and allowed to dry. Images of mScarlet and EYFP fluorescence were acquired using a Zeiss LSM 700 confocal microscope with a 20X magnification objective. Fluorescence was quantified using ImageJ (Schindelin, et al., 2012) by marking a region containing the somatic cytoplasm using the EYFP fluorescence and then measuring the average pixel intensity in the red imaging channel.

### Histology, imaging, and quantification

Mice were deeply anesthetized using pentobarbital (130□mg□per□kg, intraperitoneally) and then transcardially perfused with ice-cold PBS (pH□7.4, 10 ml) followed by 4% paraformaldehyde (PFA, 10 ml) solution. Heads were removed and post-fixed overnight at 4□°C in 4% PFA. Then, brains were extracted and transferred to 30% sucrose solution for at least 24□h. Coronal sections (40□μm) were acquired using a microtome (Leica Microsystems) and stained with a nucleic acid dye (4,6-diamidino-2-phenylindole (DAPI), 1:10,000). Slices were then mounted on gelatin-coated slides, dehydrated, and embedded in DABCO mounting medium (Sigma). Slices were imaged using a VS120 microscope (Olympus), at 10x magnification with two channels: 1) DAPI, to identify brain structures, the corresponding anterior-posterior coordinates and sites of lesions created by the optic fiber. 2) Either Cy3 (mScarlet - eOPN3 mice) or FITC (eYFP - control mice), to measure expression levels in cells and projections. The resulting images were then analyzed using ImageJ to measure the fluorescence of DAPI and additional fluorophores within specific target regions. For each slice, a rectangle outlining the target site was defined and copied to the contralateral (non-expressing) hemisphere. Mean fluorescence values were measured separately for each channel and compared between hemispheres, demonstrating differences in fluorophore expression but not in DAPI staining. Imaging acquisition parameters and the ensuing analysis pipeline were kept constant across mice to allow comparison between the eOPN3 and the control groups.

### Cell culture and live-cell cAMP assay

Optical activation and G protein coupling of mosOPN3-mScarlett and chimeric GPCR constructs was tested in HEK293T cells using a live cell assay (Ballister, et al., 2018). Briefly, GPCR constructs were subcloned into pcDNA3.1 (ThermoFisher). HEK293T cells were incubated at 37°C (5% CO_2_) in DMEM containing 4500 mg/L glucose, L-glutamine, (Sigma) with penicillin (100 U/ml), streptomycin (100 μg/ml), and 10% FBS. For transfection, cells were seeded into solid white 96-well plates (Greiner) coated with poly-L-Lysine (Sigma Aldrich) and transfected with Lipofectamine 2000 (ThermoFisher) together with individual G protein chimera (GsX) and Glo22F (Promega). Cells were incubated for 24h at 37°C, 5% CO2 and, subsequently, in L-15 media (without phenol-red, with L-glutamine, 1% FBS, penicillin, streptomycin (100 μg/ml)) and 9-cis retinal (10 μM) and beetle luciferin (2 mM in 10 mM HEPES pH 6.9) for 1 h at RT. Cells were kept in the dark throughout the entire time. Baseline luminescence was measured 3 times and opto-GPCR activation was then induced by illuminating cells for 1s with an LED plate (530 nm, 5.5 μW·mm^−2^, Phlox Corp.) Changes in cAMP levels were measured over time using GloSensor luminescence. For the assay quantification each technical repeat was normalized to its pre-light baseline.

### Slice culture preparation and transgene delivery

Organotypic hippocampal slices were prepared from Wistar rats at post-natal day 5-7 as described (Gee, et al., 2017). Briefly, dissected hippocampi were cut into 400 μm slices with a tissue chopper and placed on a porous membrane (Millicell CM, Millipore). Cultures were maintained at 37 °C, 5% CO2 in a medium containing 80% MEM (Sigma M7278), 20% heat-inactivated horse serum (Sigma H1138) supplemented with 1 mM L-glutamine, 0.00125% ascorbic acid, 0.01 mg/ml insulin, 1.44 mM CaCl_2_, 2 mM MgSO_4_ and 13 mM D-glucose. No antibiotics were added to the culture medium.

For transgene delivery in organotypic slices, individual CA3 pyramidal cells were transfected by single-cell electroporation between DIV 15-20 as previously described (Wiegert, et al., 2017). The plasmids pAAV-CKIIα(0.4)-eOPN3-mScarlet, pCI-hSyn-mCerulean, CAG-GIRK2-1 and pGP-AAV-hSyn-jGCaMP7f-WPRE were all diluted to 50 ng/μl in K-gluconate-based solution consisting of (in mM): 135 K-gluconate, 10 HEPES, 0.2 EGTA, 4 Na_2_-ATP, 0.4 Na-GTP, 4 MgCl_2_, 3 ascorbate, 10 Na_2_- phosphocreatine, pH 7.2, 295 mOsm/kg. An Axoporator 800A (Molecular Devices) was used to deliver 25 hyperpolarizing pulses (−12 V, 0.5 ms) at 50 Hz. During electroporation slices were maintained in pre-warmed (37° C) HEPES-buffered solution in (mM): 145 NaCl, 10 HEPES, 25 D-glucose, 2.5 KCl, 1 MgCl_2_ and 2 CaCl_2_ (pH 7.4, sterile filtered).

For targeted viral vector-based transduction of organotypic hippocampal slice cultures (Wiegert, et al., 2017), adeno-associated viral particles encoding AAV2/9.CamKIIα(0.4).eOPN3-mScarlet were pressure injected (20 PSI/2-2.5 bar, 50 ms duration) using a Picospritzer III (Parker) under visual control (oblique illumination) into CA3 *stratum pyramidale* between DIV 2-5. Slice cultures were then maintained in the incubator for 2-3 weeks allowing for virus payload expression.

### Electrophysiology in cultured neurons

Whole-cell patch clamp recordings in dissociated cultures were performed under visual control using differential interference contrast infrared (DIC-IR) illumination on an Olympus IX-71 microscope equipped with a monochrome scientific CMOS camera (Andor Neo). Borosilicate glass pipettes (Sutter Instrument BF100-58-10) with resistances ranging from 3–7 MΩ were pulled using a laser micropipette puller (Sutter Instrument Model P-2000). For hippocampal neuron cultures, electrophysiological recordings from neurons were obtained in Tyrode’s medium ([mM] 150 NaCl, 4 KCl, 2 MgCl_2_, 2 CaCl_2_, 10 D-glucose, 10 HEPES; 320 mOsm; pH adjusted to 7.35 with NaOH). The recording chamber was perfused at 0.5 ml min^−1^ and maintained at 29°C or 23°C (Fig. S4A). Pipettes were filled using a potassium gluconate-based intracellular solution ([mM] 135 K-gluconate, 4 KCl, 2 NaCl, 10 HEPES, 4 EGTA, 4 MgATP, 0.3 NaGTP; 280 mOsm kg^−1^; pH adjusted to 7.3 with KOH). Whole-cell voltage clamp recordings were performed using a MultiClamp 700B amplifier, filtered at 8 kHz and digitized at 20 kHz using a Digidata 1440A digitizer (Molecular Devices). Light was delivered using a Lumencor SpecraX light engine, using band-pass filters at 445/20, 475/28, 512/25, 572/35 and 632/22 nm (peak wavelength/bandwidth). Photon flux was calibrated to be similar for all five wavelengths at the sample plane to allow comparison of activation efficiency. Remaining photon flux differences were less than 6%.

Whole-cell recordings in autaptic neurons were performed on an Olympus IX73 microscope using a Multiclamp 700B amplifier (Molecular Devices) under control of Clampex 10 (Molecular Devices). Data was acquired at 10 kHz and filtered at 3 kHz. Extracellular solution contained (in mM): 140 NaCl, 2.4 KCl, 10 HEPES, 10 glucose, 2 CaCl_2_, and 4 MgCl_2_ (pH adjusted to 7.3 with NaOH, 300 mOsm). Internal solution contained the following (in mM): 136 KCl, 17.8 HEPES, 1 EGTA, 0.6 MgCl_2_, 4 MgATP, 0.3 Na_2_GTP, 12 Na_2_ phosphocreatine, 50 U ml^−1^ phosphocreatine kinase (300 mOsm); pH adjusted to 7.3 with KOH. Fluorescence light from a TTL-controlled LED system (pE4000, CoolLED) was filtered using single band-pass filters (AHF F66-415), coupled into the back port of the microscope by a liquid light guide, and delivered through an Olympus UPLSAPO 20×, 0.75 NA objective. Membrane potential was set to −70 mV, and series resistance and capacitance were compensated by 70%. To obtain strong GIRK currents, cells were voltage clamp briefly to −50 mV for the light flash only, while EPSCs were recorded at −70 mV. Synaptic transmitter release was elicited by 1 ms depolarization to 0 mV, causing an unclamped AP in the axon. Baclofen and SCH23390 were applied via a rapid perfusion system (Rost, et al., 2010). Pertussis toxin was applied to the cultures 24 h before the recordings, at a concentration of 0.5 μg ml^−1^. Cells were excluded from the analysis of the paired-pulse ratio if eOPN3 activation completely abolished the first EPSC, and mEPSCs were not analyzed when noise-events detected by an inverted template occurred at >1 Hz, as previously described (Rost, et al., 2015).

### Slice culture electrophysiology and two-photon microscopy

To characterize the effects of eOPN3-activation on neuronal cell parameters, targeted whole-cell recordings of transfected CA3 pyramidal neurons were performed at room temperature (21-23°C) under visual guidance using a BX 51WI microscope (Olympus) and a Multiclamp 700B amplifier (Molecular Devices) controlled by either Ephus (Suter, et al., 2010) or WaveSurfer software (https://wavesurfer.janelia.org/), both written in MATLAB. Patch pipettes with a tip resistance of 3-4 MΩ were filled with (in mM): 135 K-gluconate, 4 MgCl_2_, 4 Na_2_-ATP, 0.4 Na-GTP, 10 Na_2_-phosphocreatine, 3 ascorbate, 0.2 EGTA, and 10 HEPES (pH 7.2). Artificial cerebrospinal fluid (ACSF) consisted of (in mM): 135 NaCl, 2.5 KCl, 4 CaCl_2_, 4 MgCl_2_, 10 Na-HEPES, 12.5 D-glucose, 1.25 NaH_2_PO_4_ (pH 7.4). To block synaptic transmission, 10 μM CPPene, 10 μM NBQX, and 100 μM picrotoxin (Tocris, Bristol, UK) were added to the recording solution. Measurements were corrected for a liquid junction potential of −14 mV.

In dual patch-clamp experiments, we recorded from pairs of synaptically connected CA3 pyramidal cells expressing eOPN3 and non-expressing CA1 pyramidal cells. CA3 pyramidal neurons were stimulated in current clamp to elicit 2 action potentials (40 ms Inter Stimulus Interval, 0.2 Hz) by brief somatic current injection (2 - 3 ms, 3 - 4 nA) in the absence of synaptic blockers while recording EPSCs by holding the CA1 cell at −60 mV in voltage clamp mode. For extracellular stimulation, afferent Schaffer collateral axons were stimulated (0.2 ms, 20-70 μA every 10 s) with a monopolar glass electrode connected to a stimulus isolator (IS4 stimulator, Scientific Devices). For train stimulation, 10 pulses were delivered every 40 ms. Access resistance of the recorded non-transfected CA1 neuron was continuously monitored and recordings above 20 MΩ and/or with a drift > 30% were discarded. A 16 channel pE-4000 LED light engine (CoolLED, Andover, UK) was used for epifluorescence excitation and light activation of eOPN3 (500ms, 525 nm, 1 mW mm^−2^). Light intensity was measured in the object plane with a 1918 R power meter equipped with a calibrated 818 ST2 UV/D detector (Newport, Irvine CA) and divided by the illuminated field of the Olympus LUMPLFLN 60XW objective (0.134 mm^2^) or of the Olympus LUMPLFLN 40XW objective (0.322 mm^2^) in dual-patch and extracellular stimulation experiments. All the electrophysiological synaptic measurements in organotypic hippocampal slice cultures were performed at 33 ± 1 °C.

For the eOPN3 two-photon stimulation experiments, a custom-built two-photon imaging setup was used based on an Olympus BX51WI microscope controlled by ScanImage 2017b (Vidrio Technologies). Electrophysiological recordings were acquired using a Multiclamp 700B amplifier controlled by the WaveSurfer software written in MATLAB (https://wavesurfer.janelia.org/). A tunable, pulsed Ti:Sapphire laser (MaiTai DeepSee, Spectra Physics) controlled by an electro-optic modulator (350-80, Conoptics) tuned to 1040 nm was used to excite the mScarlet-labeled eOPN3. Red fluorescence was detected through the objective (LUMPLFLN 60XW, 60x, 1.0 NA, Olympus) and through the oil immersion condenser (numerical aperture 1.4, Olympus) by photomultiplier tubes (H7422P-40SEL, Hamamatsu). 560 DXCR dichroic mirrors and 525/50 and 607/70 emission filters (Chroma Technology) were used to separate green and red fluorescence. Excitation light was blocked by short-pass filters (ET700SP-2P, Chroma). In addition, the forward-scattered IR laser light was collected through the condenser, spatially filtered by a Dodt contrast tube (Luigs&Neumann) attached to the trans-illumination port of the microscope and detected with a photodiode connected to a detection channel of the laser scanning microscope. This generated an IR-scanning gradient contrast image (IR-SGC) synchronized with the fluorescence images. (Wimmer, Nevian et al. 2004). This approach was used for targeted patch-clamp recordings avoiding prior activation of the ultrasensitive eOPN3 with epifluorescence illumination. The two-photon laser scanning pattern used for stimulation was either a spiral scan with a repetition rate of 500 Hz above the soma (2 ms/spiral, 250 cycles, 500 ms total duration) or standard raster scans at 1.09 Hz over the somatodendritic compartment (FOV=106*106 μm, 512×512 pixels, 1.8 ms/line, 5 frames, 4.6 s total duration). The laser wavelengths used for stimulation were 800 nm, 860 nm, 930 nm, 980 nm and 1040 nm, all at 30 mW, measured at the back focal aperture of the objective. Wide field illumination at 525 nm (10 mW/mm^2^) was done with a 16 channel pE-4000 LED light engine (CoolLED, Andover, UK) for 500 ms. An additional set of experiments was performed on a second custom-modified two-photon imaging setup (DF-Scope, Sutter) based on an Olympus BX51WI microscope controlled by ScanImage 2017b (Vidrio Technologies) and equipped with an Ytterbium-doped 1070-nm pulsed fiber laser (Fidelity-2, Coherent) for far infrared stimulation. Electrophysiological recordings were performed using a Double IPA integrated patch amplifier controlled with SutterPatch software (Sutter Instrument).

The same microscope was used to acquire images of eOPN3-expressing CA3 cells co-transfected with the cyan cell-filler fluorophore mCerulean (Rizzo, et al., 2004) and their projecting axons in *stratum radiatum* of CA1. The 1070-nm laser was used to excite fluorescence of mScarlet-labeled eOPN3. mCerulean was excited by a pulsed Ti:Sa laser (Vision-S, Coherent) tuned to 810 nm. Laser power was controlled by electro-optic modulators (350-80, Conoptics). Red and cyan fluorescence were detected through the objective (Olympus LUMPLFLN 60XW, 1.0 NA, or Leica HC FLUOTAR L 25x/0.95 W VISIR) and through the oil immersion condenser (numerical aperture 1.4, Olympus) by GaAsP photomultiplier tubes (Hamamatsu, H11706-40). Dichroic mirrors (560 DXCR, Chroma Technology) and emission filters (ET525/70m-2P, ET605/70m-2P, Chroma Technology) were used to separate cyan and red fluorescence. Excitation light was blocked by short-pass filters (ET700SP-2P, Chroma Technology). All electrophysiology recordings were analyzed using custom written scripts in MATLAB except for recordings acquired with the Double IPA integrated patch amplifier, which were analyzed with the SutterPatch software.

For presynaptic Ca^2+^ imaging experiments, a custom-modified version of ScanImage 3.8 (Pologruto et al., 2003) was used to allow user-defined arbitrary line scans. jGCaMP7f was excited at 960 nm. Similar to the two-photon stimulation experiments, targeted patch-clamp recordings were achieved using IR-scanning gradient contrast image (IR-SGC) synchronized with the fluorescence images. Single action potentials were triggered by brief somatic current injection (2 - 3 ms, 3 - 4 nA) in the absence of synaptic blockers while monitoring fluorescent transients at single Schaffer collateral terminals in CA1 (70-80 trials on average at 0.1 Hz). User-defined circular scans at 500 Hz across the bouton were used to repeatedly sample the fluorescent changes. During each trial (3 s), laser exposure was restricted to the periods of expected Ca^2+^ response (~1.3 s) to minimize bleaching. To activate eOPN3 selectively at the terminals, we used a fiber-coupled LED (400 mm fiber, NA 0.39, M118L02, ThorLabs) to deliver 500 ms green light pulses (λ = 530 nm, 83μW at the fiber tip) 1 s prior to the onset of electrical stimulation. During the LED pulses, upper and lower PMTs were protected by TTL triggered shutters (NS45B, Uniblitz). GIRK channels were blocked by SCH 23390 (10 μM, Tocris, Bristol, UK) throughout the entire experiment to exclude hyperpolarization-mediated effects on action potential propagation and presynaptic Ca^2+^ influx.

The photon shot-noise subtracted relative change in jGCaMP7f fluorescence (ΔF/F_0_) was measured by using a template-based fitting algorithm. The characteristic fluorescence time constant was extracted for every bouton by fitting a double exponential function (τ_rise_, τ_decay_) to the average jGCaMP7f signal. To estimate the Ca^2+^ transient amplitude for every trial, we fitted the bouton-specific template to every response, amplitude being the only free parameter. Response amplitude was defined as the value of the fit function at its maximum.

### *In vivo* electrophysiological recordings

8-9 weeks old male C75/Bl6 mice were pressure injected (Picospritzer III; Parker) bilaterally into LGN (AP: − 2.2 mm; ML: +/− 2.3 mm; DV: −3.1 mm) at 50 nl/min with 200 nl adeno-associated viral particles encoding eOPN3 (AAV2/5.CKIIa(0.4).eOPN3-mScarlet) diluted to 2.5 × 10^12^ viral genomes per ml using a pulled glass capillary. Following 5-6 weeks of recovery, mice underwent 3-4 head fixation habituation sessions starting with 15 min and gradually increasing to 25 min. 7-12 weeks after virus injection, craniotomies were performed bilaterally to provide access to V1 spanning from −2.3 mm to −4.7 mm in the anterior posterior direction and 2 mm at its widest part (at AP: −3.8 mm) from +/− 1.3 mm to +/− 3.3 mm along the medio-lateral axis. Craniotomies were covered with Kwik-Cast (WPI Inc) to protect the brain surface from mechanical impact, dehydration, and light exposure between the silicon probe recording sessions.

For the electrophysiological recordings, two 4-shank, 128 channel silicon microprobes (128DN; 4 shanks, 150 μm shank spacing, 25 μm channel spacing, 100 μm^2^ electrode area, 7 mm × 65 μm × 23 μm shank dimensions) (Yang, et al., 2020) (kindly provided by Dr. S. Masmanidis, UCLA) were inserted bilaterally in the V1 at a depth of approximately 1 mm, with an insertion speed of 100 μm/min. Before each recording session, silicon probe recording sites were electroplated in a PEDOT solution to an impedance of ~100 kOhm. Each silicon probe was connected to an RHD2000 chip-based 128 channel amplifier board (Intan Technologies). Broadband (0.1 Hz-7.5 kHz) signals were acquired at 30 kHz. Signals were digitized at 16 bit and transmitted to an OpenEphys recording controller (OEPS).

Raw data were processed to detect spikes and extract single-unit activity. Briefly, the wide-band signals were band-pass filtered (0.6 kHz-6 kHz), spatially whitened across channels and thresholded for isolation of putative spikes. Clustering was performed using template matching implemented in Kilosort2 (Pachitariu, et al., 2016) and computed cluster metrics were used to pre-select units for later manual curation using custom-written software.

For the optogenetic inhibition of LGN axons, the silicon probe inserted in one of the two craniotomies was coupled with a 200 μm 0.5 NA optic fiber (Thorlabs, FP200URT), placed between the two middle shanks and at ~300 μm above the top-most channel of the silicon probe, thus the optic fiber remained just outside the surface of the cortex during the recordings. This fiber was coupled with a 525 nm LED (PlexBright, Plexon), controlled using a Cyclops 3.6 LED driver and a custom Teensy3.2-based stimulation system, calibrated to deliver ~2 mW of light at the tip of the fiber.

Following a long baseline period, the paradigm used to investigate the effect of eOPN3 on the synaptic vesicle release *in vivo* consisted of 31 presentations of a visual stimulus every 30 seconds. The 10 first trials were used to establish the baseline of the visual response and the 11th trial was coupled with optogenetic stimulation, starting 1 second before the visual stimulation and lasting for a total of 30 sec. Each visual stimulus presentation trial consisted of 8 repeats of a 500 ms visual drifting grating presentations in the cardinal and intercardinal directions. The stimuli were presented on a 23.5” monitor placed 20 cm centrally in front of the mouse, so that the monitor was visible to both eyes. The stimulus presentation was controlled using a custom-written Python program and utilized PsychoPy3.0. For the accurate detection of the stimulus onset to allow for alignment with electrophysiological data, a photodetector was mounted in one corner of the monitor. The mouse was gradually habituated to head-fixation over multiple sessions and was running freely on a horizontal wheel. Each mouse was recorded for 1 or 2 identical sessions on different days and data were pooled for the subsequent analyses. Recording sessions in which no units showed visual stimulus-evoked activity were excluded from the analysis.

For visual stimulus response characterization, the spike rates were calculated in 50 ms bins. Each unit’s activity was normalized to the average firing rate in the 15 s prior to stimulus presentation during the baseline period. The baseline period in Fig. S7D was defined as the activity during the two trials before eOPN3 activation. For clarity, the peristimulus time histograms shown in Fig. S7E were low pass filtered using a Gaussian function (window: 250 ms, σ = 100 ms). The recovery time constant shown in Fig. S7F was calculated by fitting the post eOPN3 activation visual stimulus response to f(t) = 1-a*exp(-t/tau), with the effect size (a) and recovery time constant (tau) as free parameters.

### *In vivo* optogenetic silencing of the nigrostriatal pathway

AAV vectors encoding a Cre-dependent eOPN3-mScarlet transgene (AAV2/1&2.hSyn.SIO-eOPN3-mScarlet; 6E12 viral genomes / ml) or eYFP (AAV2/1&2.EF1a.DIO.eYFP; 2E13 viral genomes / ml) were unilaterally injected into the substantia nigra (AP: − 3.5 mm, ML: + or − 1.4 mm DV: − 4.25 mm; 500 nl per mouse). Optical fibers (200 μm diameter, NA 0.5) were unilaterally implanted above the ipsilateral dorsomedial striatum (AP: + 0.6 mm, ML: + or – 1.5 mm DV: − 2.1 mm). Left and right implanted mice were counterbalanced among the eOPN3 and control groups. Mice were allowed to recover for 6-9 weeks to allow for viral expression. Following recovery, mice underwent a single 10-minute habituation session, to habituate to handling, patch cord attachment and the open field arena. In experimental sessions, we attached individual mice to a patch cord and video recorded their free locomotion continuously in the open field under near-infrared illumination.

To measure eOPN3 induced bias in locomotion, we video recorded the free locomotion of single mice in an open field arena (50 × 50 × 50 cm) continuously over 30 minutes. After a 10-minute baseline no-light period, we delivered 500 ms light pulses (540 nm, 10 mW at the fiber tip), at 0.1 Hz for 10 minutes, followed by an additional 10-minute no-light period. Offline video processing and mouse tracking was done using Deep Lab Cut (DLC; (Mathis, et al., 2018)). Briefly, we trained DLC to detect 6 features on the mouse body (nose, head center, left and right ears, center of mass, tail) and 3 bottom corners of the arena. X-Y coordinates of each feature were then further processed to complete missing or noisy values (high amplitude and short duration changes in X or Y dynamics) using linear interpolation (*interp1*) of data from neighboring frames. This was followed by a low pass filtering of the signals (*malowess*, with 50 points span and of linear order). Finally, a pixel to cm conversion was done based on the video-detected arena features and its physical measurements. A linear fit to the nose, head, center and tail features defined the mouse angle with respect to the south arena wall at each frame. Following its dynamics over the session, we identified direction shifts as a direction change in angle that exceeds 20° and 1 second. To achieve a comparable measurement between right- and left-hemisphere injected mice, we measured motion in the ipsilateral direction as positive and contralateral motion as negative from the cumulative track of angle. The net angle gain was calculated as the sum of ipsilateral and contralateral angle gained over each time bin (1- or 10-minute bins as indicated). For each time bin we then calculated a rotation index, based on angle gains, as follows:

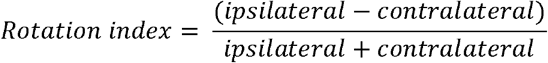

For each mouse, rotation index scores were calculated for two complete sessions on different days. Individual scores were plotted for each mouse against the expression levels measured in that mouse (see section: Histology, imaging, and quantification). Results were then averaged across individual sessions, and used for all statistical comparisons, and linear regressions analysis. Mouse positions and velocities were measured by the center feature.

## Author contributions

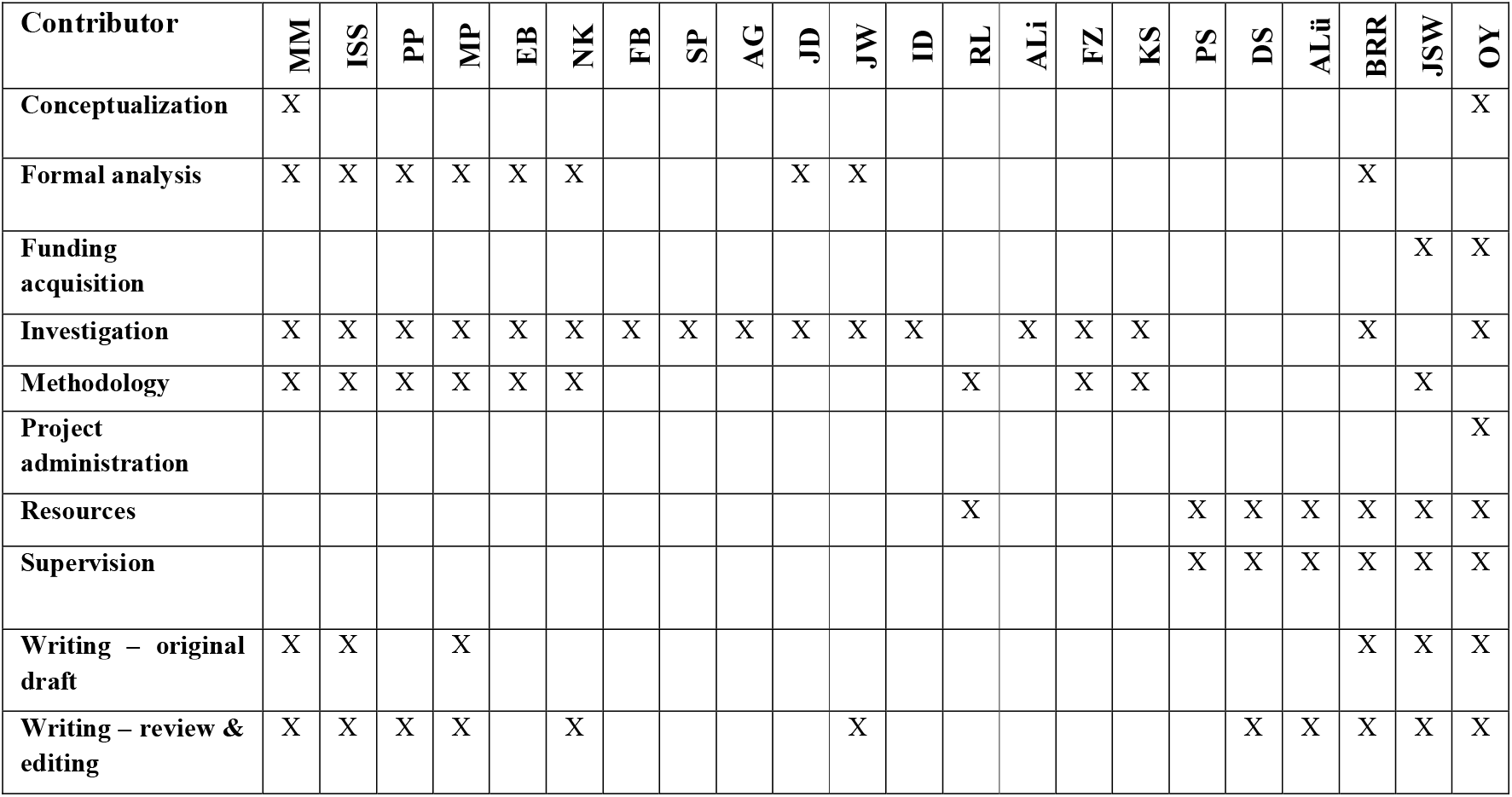

## Acknowledgements

We thank the members of the Yizhar, Wiegert, Soba and Schmitz labs for ideas, criticism, and discussions throughout this project. We thank Bryan Copits, Michael Bruchas and Robert Gereau for insightful comments on the manuscript. We would also like to thank Thomas Oertner for generous sharing of equipment and Eitan Reuveny for the GIRK expression plasmids. This work was supported by funding from the European Research Commission (ERC CoG PrefrontalMap 819496 to OY, ERC StG LIFE synapses to 714762 JSW), the Israel Science Foundation (COEX 3131/20), the Adelis Foundation, the Candice Appleton Family Trust (to OY), the Achar Research Fellow Chair in Electrophysiology (to JD), and the German Research Foundation (DFG FOR2419 to JSW and DFG SPP 1926 jointly to OY, PS, and JSW).

## Supplemental figures

**Figure S1.**
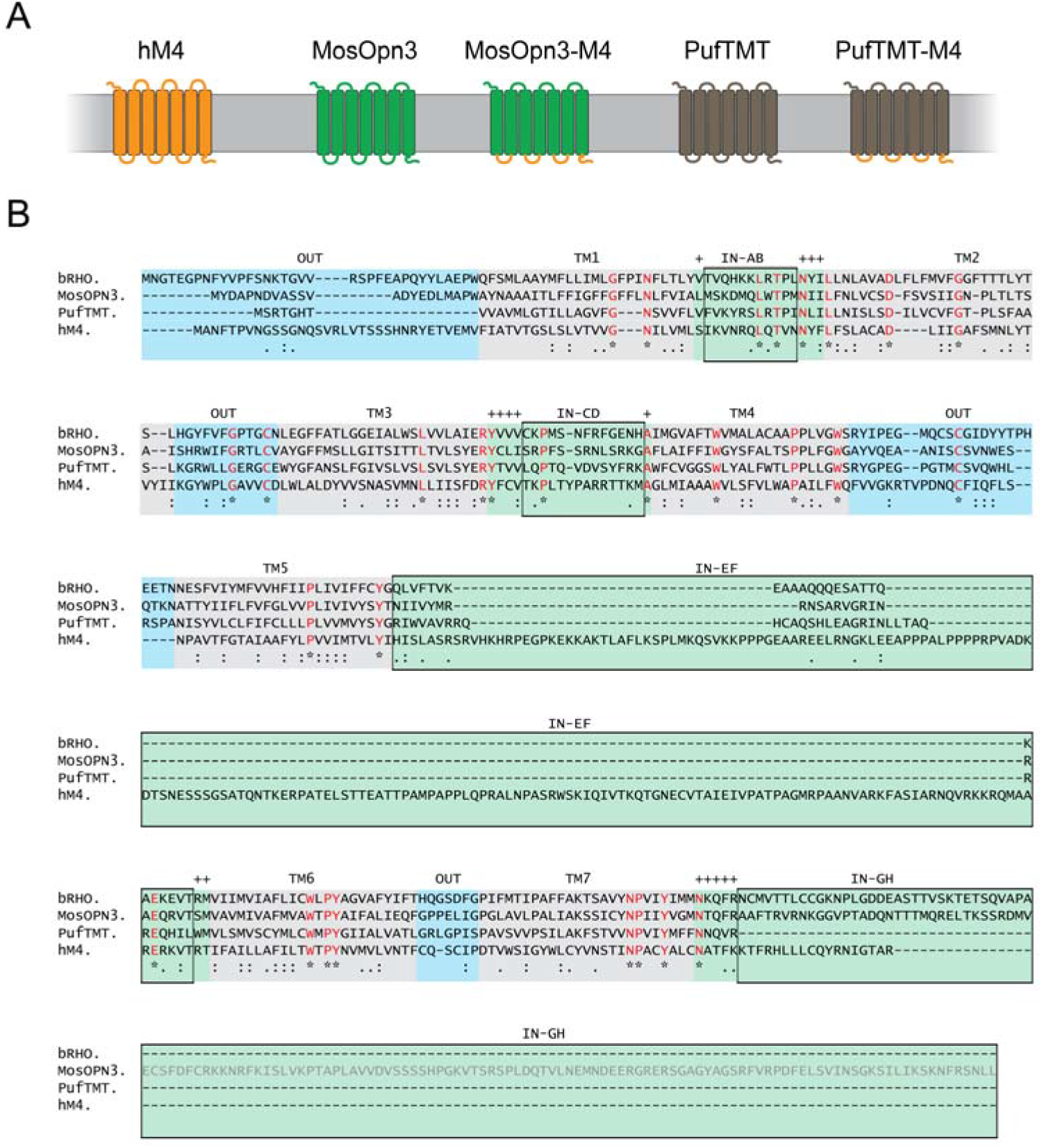
hM4 chimera design. **(A)** Schematic diagrams of chimeric proteins comprising transmembrane and extracellular domains from the bistable mosquito OPN3 opsin (OPN3, GenBank: AB753162.1) or the teleost multiple tissue opsin from pufferfish (PufTMT, GenBank: AB753163.1) and intracellular domains of the human muscarinic receptor 4 (hM4, GenBank: NM_000741). **(B)** Multiple sequence alignment (Edgar, 2004) of the amino acid sequences of visual and non-visual rhodopsins, along with hM4. Shown are sequences of the bovine rhodopsin (bRho), OPN3, PufTMT, and hM4. Intracellular domains are labeled with green background, extracellular domains are labeled with blue background and the transmembrane domains are in gray. “*” indicates an identical amino acid in all sequences in the alignment (red letters), “:” indicates conserved amino acid substitutions according to the COLOUR table (http://www.jalview.org/help/html/colourSchemes/clustal.html), and “.” indicates semi-conserved substitutions. Intracellular regions that were replaced by the hM4 sequence to create chimeric proteins are indicated by black boxes. Non-replaced amino acids within the intracellular region are indicated by a + above the column. The 99 amino acid deletion in OPN3, introduced to improve expression in neurons, is indicated by gray amino acid letters (bottom row).

**Figure S2.**
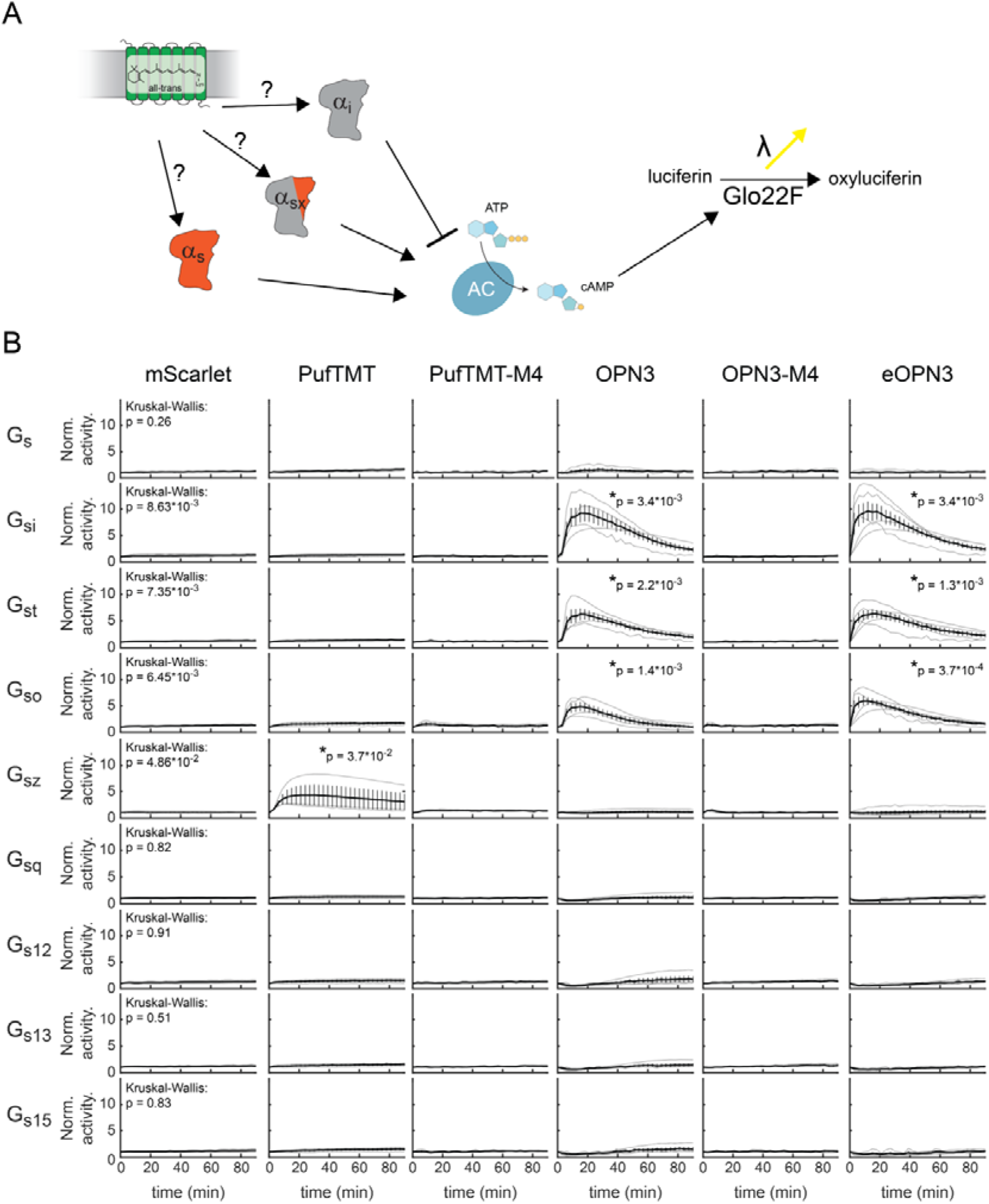
G protein activation assay. Light-dependent G protein activation by several opto-GPCR constructs, assayed in HEK293T cells. **(A)** Essay scheme. HEK293T cells are transfected with chimeras of G_α_ proteins and the G_αs_ C-terminus. cAMP levels in live cells are measured through the cAMP reporter (Glo22F). This allows for measuring cAMP levels as readout of chimera activation by the co-expressed opto-GPCR. **(B)** opto-GPCRs were activated with a green LED pulse (1s, 530nm, 5.5 μW·mm^−2^) and luminescence was measured over time. Graphs show the light-induced response, normalized to pre-activation baseline, for mScarlet (control, n = 4), PufTMT-mScarlet (n = 3), pufTMT-M4-mScarlet (n = 3), OPN3-mScarlet (n = 4), OPN3-M4-mScarlet (n = 3), and eOPN3-mScarlet (n = 5). Only OPN3-mScarlet and eOPN3-mScarlet specifically and strongly activated inhibitory G proteins (G_i_, G_t_, G_o_) in a light-dependent manner (Kruskal-Wallis tests of the maximal measured values per G protein, followed by Bonferroni-Holm corrected pairwise comparisons using Conover–Iman tests; reported p-values describe the comparison against the mScarlet control). Single trials are depicted in gray, mean ± SEM are in black.

**Figure S3.**
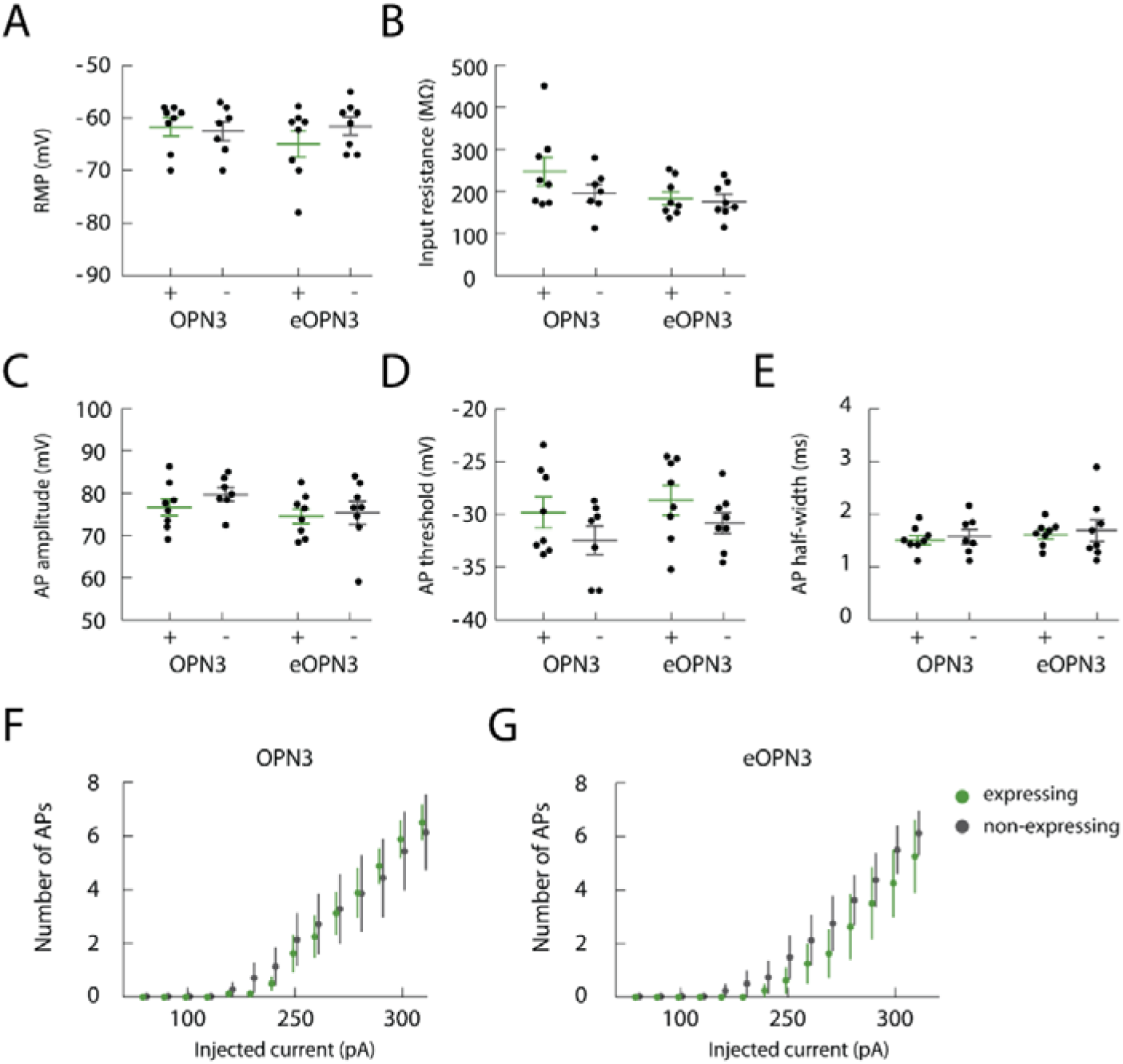
No change in the intrinsic excitability of cultured hippocampal neurons expressing OPN3-mScarlet or eOPN3-mScarlet in the absence of light. The following intrinsic properties were characterized in cultured hippocampal neurons: **(A)** resting membrane potential (RMP, OPN3 vs. ctrl: p = 0.79; eOPN3 vs. ctrl: 0.27; two-tailed Mann-Whitney tests), **(B)** membrane input resistance (OPN3 vs ctrl: p = 0.35; eOPN3 vs. ctrl: 0.82; two-tailed Mann-Whitney tests), **(C)** action potential (AP) amplitude (OPN3 vs. ctrl: p = 0.19; eOPN3 vs. ctrl: 0.57; two-tailed Mann-Whitney tests), **(D)** AP threshold (OPN3 vs. ctrl: p = 0.38; eOPN3 vs. ctrl: 0.23; two-tailed Mann-Whitney tests), and **(E)** AP half-width (OPN3 vs. ctrl: p = 0.85; eOPN3 vs. ctrl: 0.94; two-tailed Mann-Whitney tests). No differences between neurons expressing OPN3-mScarlet (n = 7) or eOPN3-mScarlet (n = 8) and neighboring non-transfected control cells (n = 7 and n = 8, respectively) were detected. **(F-G)** The number of evoked APs in response to current injection were not different in neurons expressing OPN3 or eOPN3 and non-expressing controls (p = 0.91 and 0.46, respectively; two-way repeated measures ANOVA). Plots show individual data points and average ± SEM.

**Figure S4:**
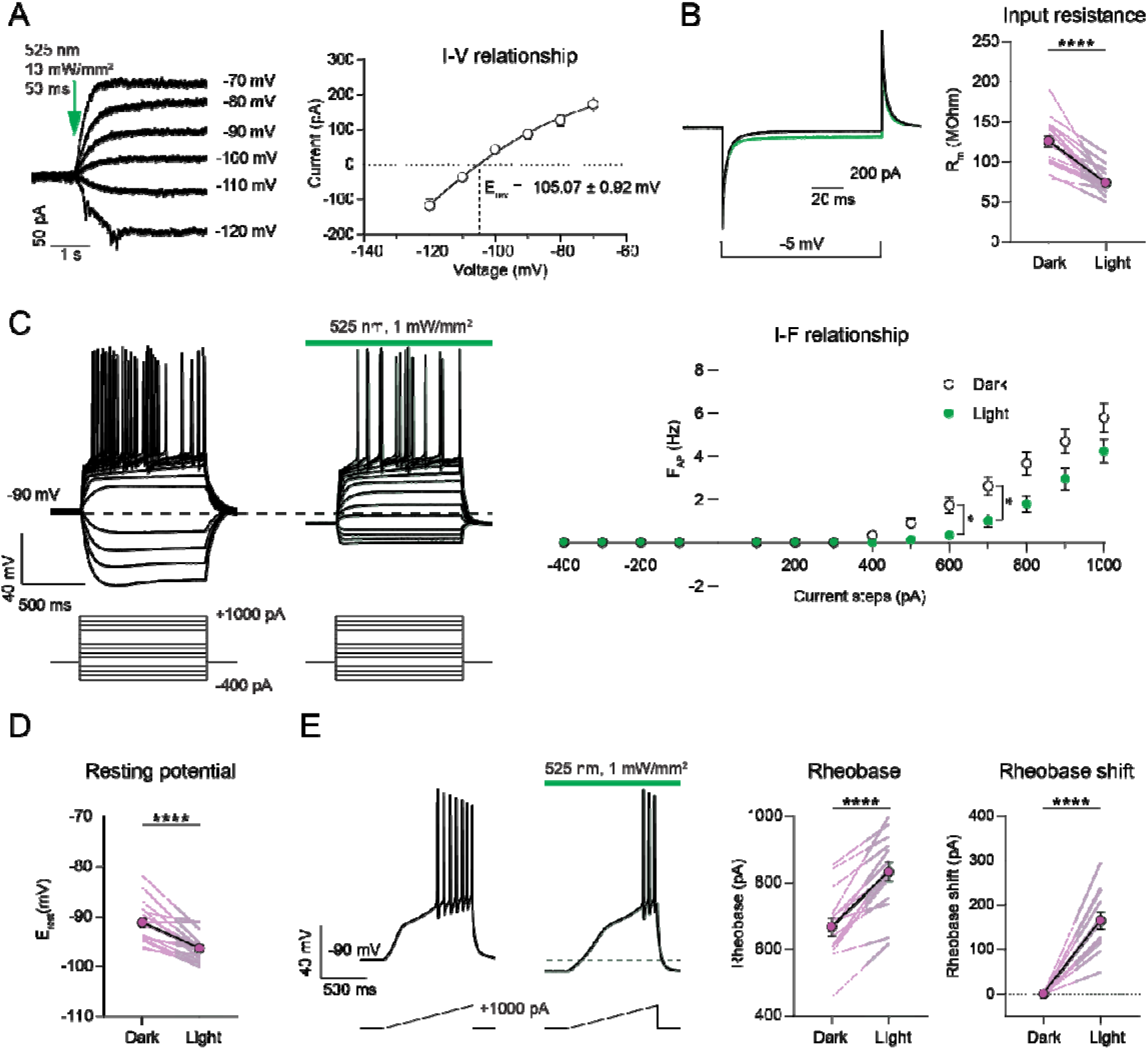
Passive and active membrane properties of eOPN3-expressing CA3 pyramidal neurons in organotypic hippocampal slices. **(A)** Light-evoked (putative GIRK) currents evoked by 50-ms green-light pulses (525 nm, 10 mW·mm^−2^) at different holding potentials, ranging from −70 to −120 mV. Values are baseline-subtracted and corrected for a liquid junction potential of −14 mV. Representative traces are shown on the *left*, quantification of the current-voltage relationship is shown on the *right* (n = 6). The photocurrent reversal potential of −105.07 ± 0.92 mV (determined with a non-linear fit) is close to the calculated K^+^ equilibrium potential of −102.5 mV. **(B)** *Left*: Representative current traces in response to a negative voltage step (−5 mV, 100 ms) in the dark (black traces) and during continuous green light (525 nm, 1 mW·mm^−2^). Note the drop of the stationary current resulting from a decreased input resistance due to increased GIRK channel conductance under illumination. *Right*: Quantification of input resistance. (Dark: 126 ± 6.79 MΩ, Light: 73 ± 3.46 MΩ, p < 0.0001, Wilcoxon-test, n = 18). **(C)** *Left*: representative voltage responses to somatic current injections ranging from −400 pA to +1000 pA in the dark and during illumination (525 nm, 1 mW·mm^−2^). *Right*: I-F plot showing decreased spike frequency in response to positive current injections, likely due to G_i/o_-mediated GIRK channel opening (p < 0.05, n = 18, two-way ANOVA with Sidak’s multiple comparisons test). **(D)** Quantification of the resting membrane potential from the current step experiments shown in C (Dark: −91.18 ± 0.96 mV; Light: −96.34 ± 0.62 mV; p < 0.0001, paired t-test, n = 18). **(E)** *Left*: representative voltage traces in response to depolarizing current ramps to assess the eOPN3-mediated rheobase shift (0 - 1000 pA). Injected current at the time of the first spike was defined as the rheobase. Green light (525 nm, 1 mW·mm^−2^) raised the rheobase of current-ramp-evoked APs. *Right*: quantification of the absolute rheobase (dark: 667.9 ± 26.79 pA, light: 832.7 ± 28.69 pA; p < 0.0001, paired t-test, n = 15) and the rheobase shift (light: 164.8 ± 19.30 pA, p < 0.0001, paired t-test, n = 15).

**Figure S5.**
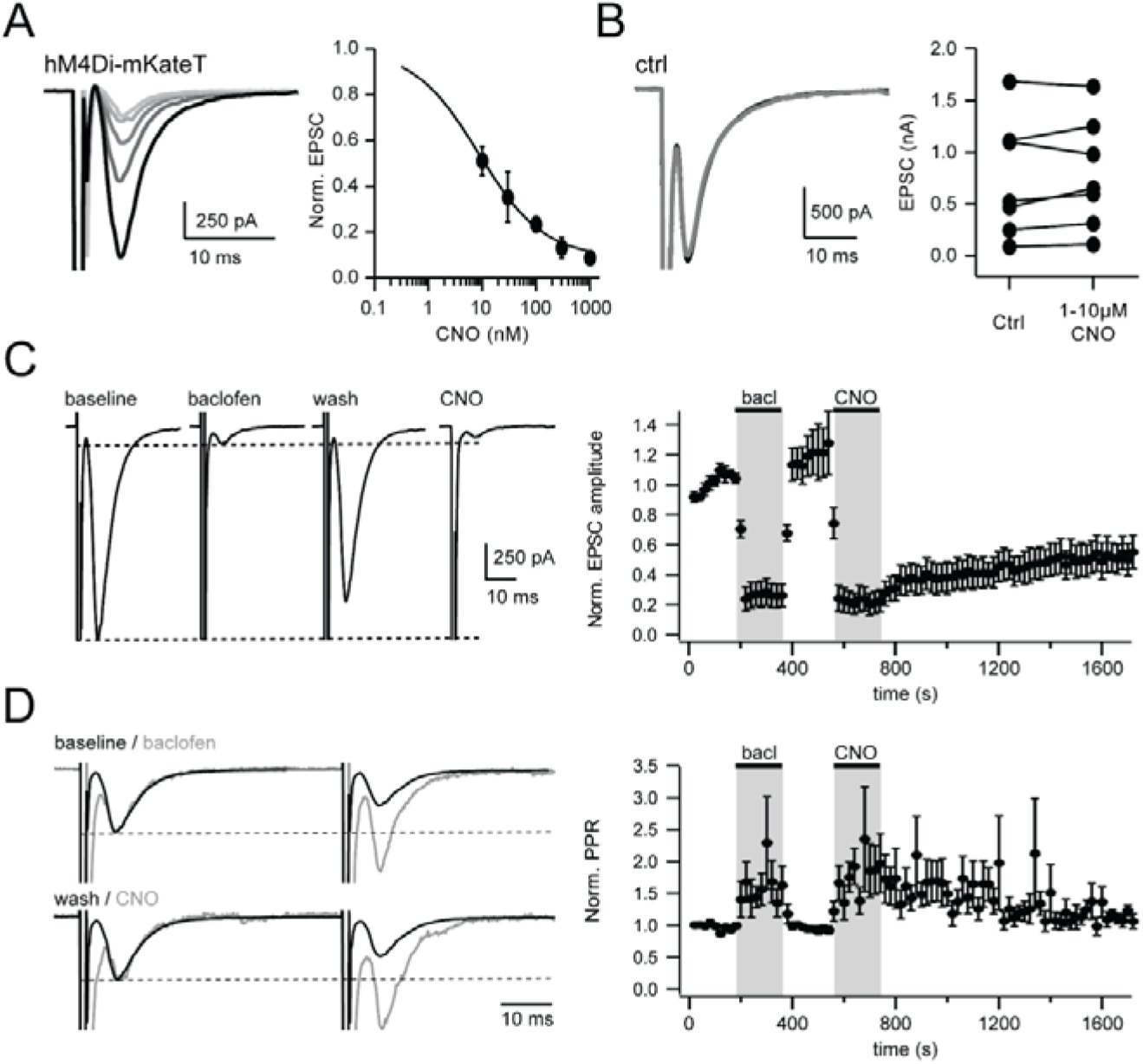
Presynaptic inhibition of neurotransmitter release by hM4Di expressed in autaptic cultures of hippocampal neurons. **(A)** Application of increasing concentrations of clozapine-N-oxide (CNO; 10, 30, 100, 300, 1000 nM, from black to light gray) leads to reduction in EPSC amplitude (IC_50_ = 8.6 nM, n = 3-12). **(B)** CNO (1-10 μM) has no effect on EPSC amplitude in neurons not expressing hM4Di. **(C-D)** Comparison of presynaptic inhibition by GABA_B_R and presynaptic inhibition by hM4Di. After 30 μM baclofen application for 90 s and washout, 100 nM CNO was added for 90 s to the same cells. Action potentials were evoked by depolarization to 0 mV for 1 ms at 0.2 Hz. Data were binned by 2, n = 5. **(C)** Both types of GPCRs suppress EPSC amplitudes to a similar extent. However, washout kinetics of CNO is dramatically slower compared to baclofen. **(D)** Increased paired-pulse ratio in response to both GABA_B_ and hM4Di receptor activation, indicating a presynaptic action. Example traces are scaled to the peak of the first EPSC under control conditions for both baclofen and CNO applications.

**Figure S6:**
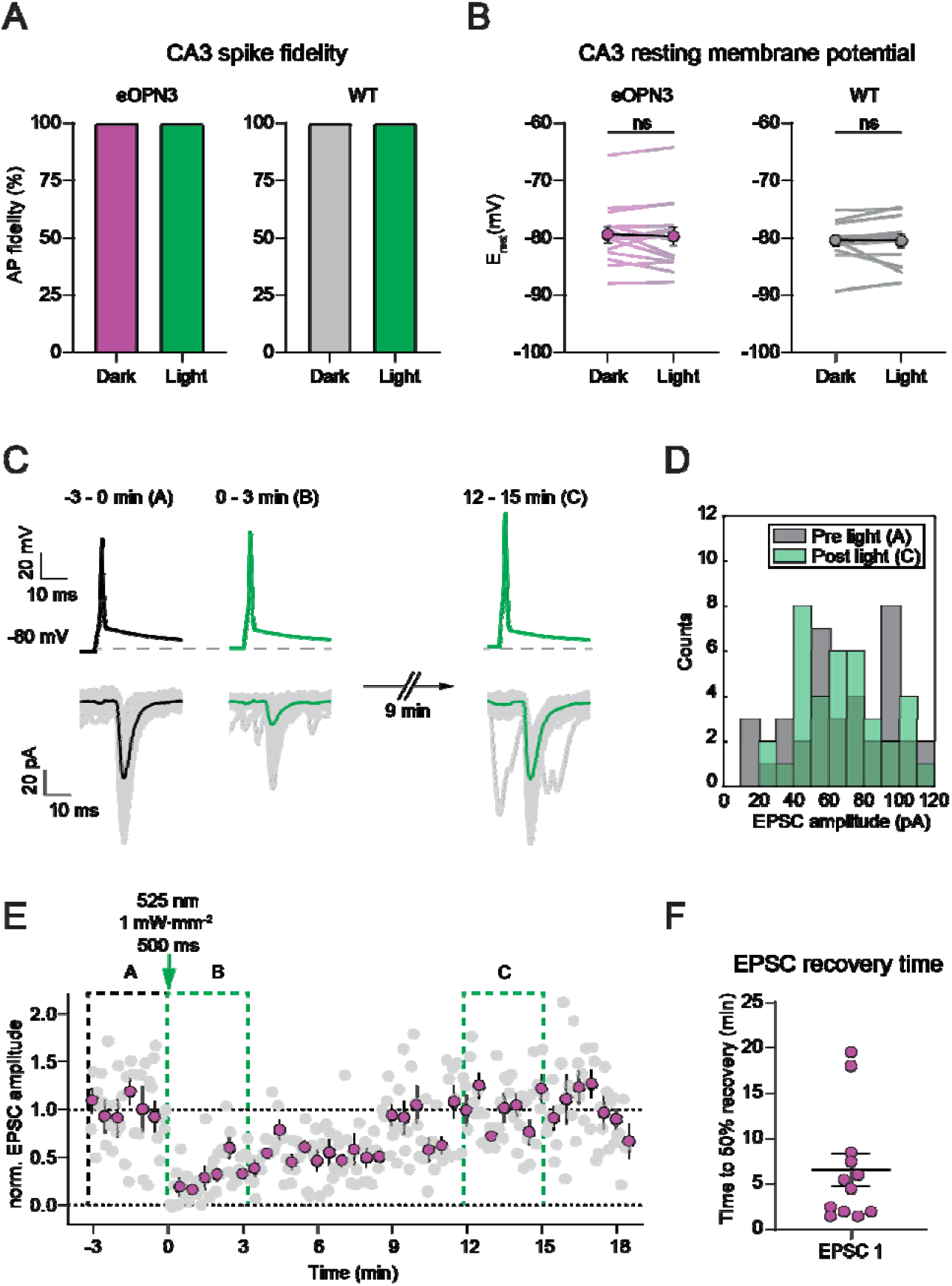
Excitability of CA3 neurons and EPSC recovery in paired-recording experiments. **(A)** Comparison of action potential success rate in CA3 in the dark and in the 30 s after light stimulation in CA1 (eOPN3 dark, eOPN3 light = 100%, n = 14; WT dark, WT light = 100%, n = 13). **(B)** Quantification of the resting membrane potential of CA3 pyramidal cells used in paired recordings in the dark and in the 30 s after light stimulation in CA1 (500 ms of 525 nm light at 1 mW·mm^−2^; eOPN3 dark: −79.41 ± 1.43, eOPN3 light: −79.71 ± 1.62, p = 0.9032, Wilcoxon test, n = 14; WT dark: −80.41 ± 0.94, WT light: −80.47 ± 1.14, p = 0.3396, Wilcoxon test, n = 13). Plots show individual data points (lines), and average (circles) ± SEM. Note absence of effects of local CA1 illumination on CA3-cell somatic properties. **(C)** Representative voltage (*top*) and current (*bottom*) traces from the example shown in E. For display purposes “pulse 2” of the paired-pulse stimulation was omitted. Note the EPSC recovery within minutes after light application. **(D)** Histogram count of peak current amplitudes of the example shown in C. **(E)** Quantification of the normalized EPSC peak amplitude shown in C (gray: individual trials, magenta: 30 s bins). **(F)** The EPSC recovery time was defined as the first 30 s-bin post light reaching at least 50% recovery of the EPSC peak amplitude compared to the average baseline EPSC peak amplitude (EPSC 1: 6.58 ± 1.78 min, mean + SEM, n = 12). Each circle represents an individual paired recording experiment.

**Figure S7:**
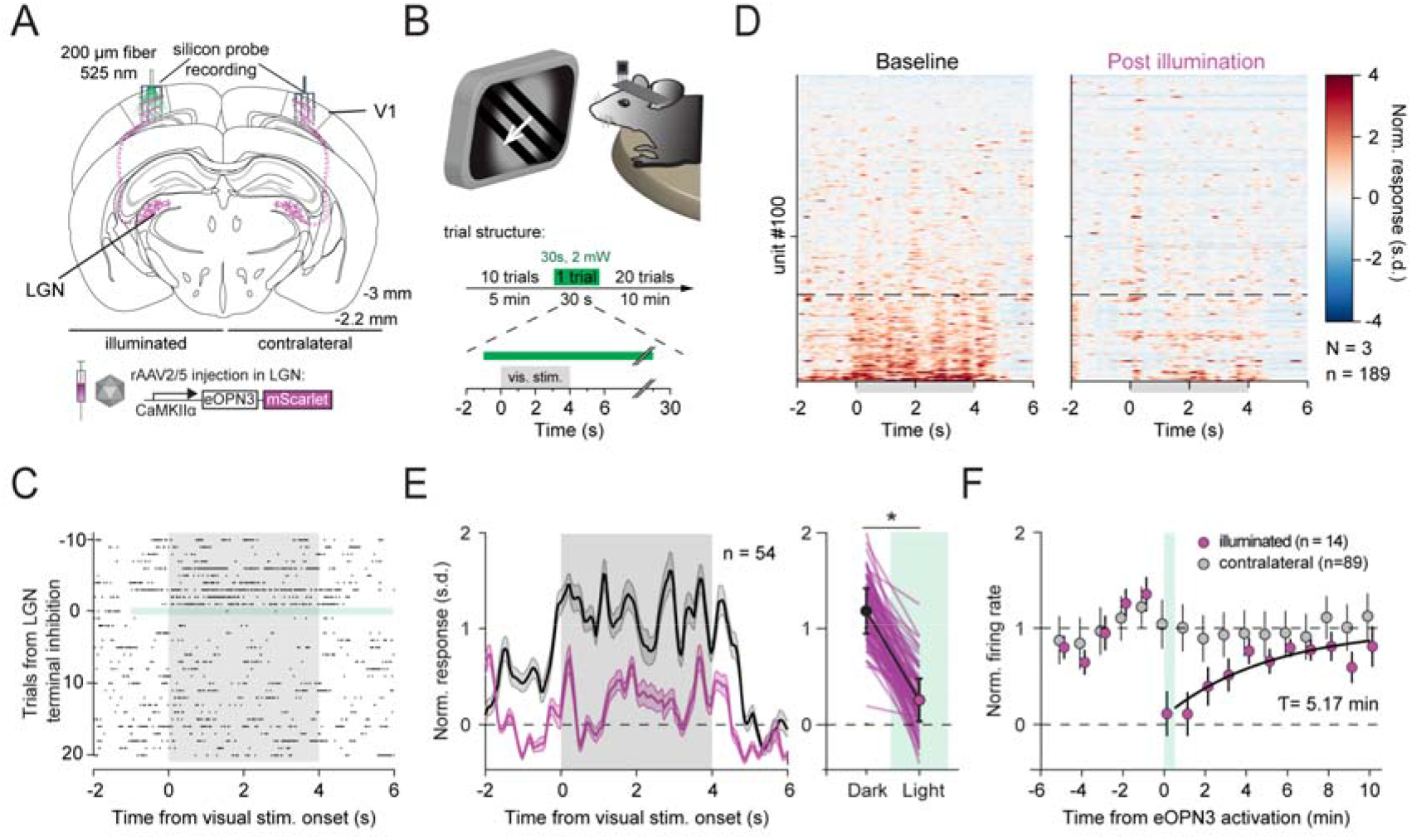
eOPN3 mediated suppression of thalamocortical inputs in awake head-fixed mice. **(A)** Schematic diagram of the investigated circuit. Lateral geniculate nucleus (LGN) neurons were bilaterally transduced with eOPN3. Acute silicon probe recordings were performed in primary visual cortex (V1) before and after unilateral illumination of LGN terminals in V1. **(B)** During silicon probe recordings, head-fixed mice were presented with a compound drifting grating stimulus (4 s duration) every 30 s for 21 trials (top). The trial structure (bottom) consisted of 10 baseline trials, followed by a single trial paired with 30 s of light delivery (525 nm at ~2 mW from a 200 μm, 0.5 NA optical fiber) to V1, and 20 post stimulus trials. **(C)** Raster plot of a representative V1 unit with reduced firing rate induced by eOPN3 activation. **(D)** Heat plot of the population response to visual stimulus presentation of all recorded units (189 units from 3 mice) on the hemisphere of eOPN3 activation before (left) and after (right) eOPN3 activation. Units were sorted by their visual stimulus presentation response magnitude during baseline condition. Units below the dashed line (n = 54) show a positive average response during the 4 s visual stimulus presentation. **(E)** *Left:* Average peristimulus time histogram of the visual stimulus responsive units (below dashed line in D). Each unit’s activity was normalized to the average firing rate in the 15 s prior to stimulus presentation during the two trials before eOPN3 activation. *Right:* Quantification of the average response during 4 s visual stimulus presentation in the two trials before (Dark) and first two trials after eOPN3 activation onset (Light). Dark: 1.17 ± 0.23, Light: 0.25 ± 0.22, *p* < 0.001, Wilcoxon test, n = 54 units. Plot shows individual units (lines), and population average (circles) ± SEM. **(F)** Kinetics of the recovery of visual stimulus response amplitude. Units from the illuminated hemisphere that showed a stimulus-evoked response reduction of at least 50% compared to the mean response in the 10 trials prior to eOPN3 activation were sub selected and their visual stimulation response recovery following the end of eOPN3 activation (magenta) was fitted with a mono-exponential function (black line). Units recorded simultaneously from the contralateral hemisphere (gray) did not change their response following ipsilateral eOPN3 activation. During the baseline and post light period, the plot shows the averages of two consecutive trials (circles) ± SEM.

**Figure S8:**
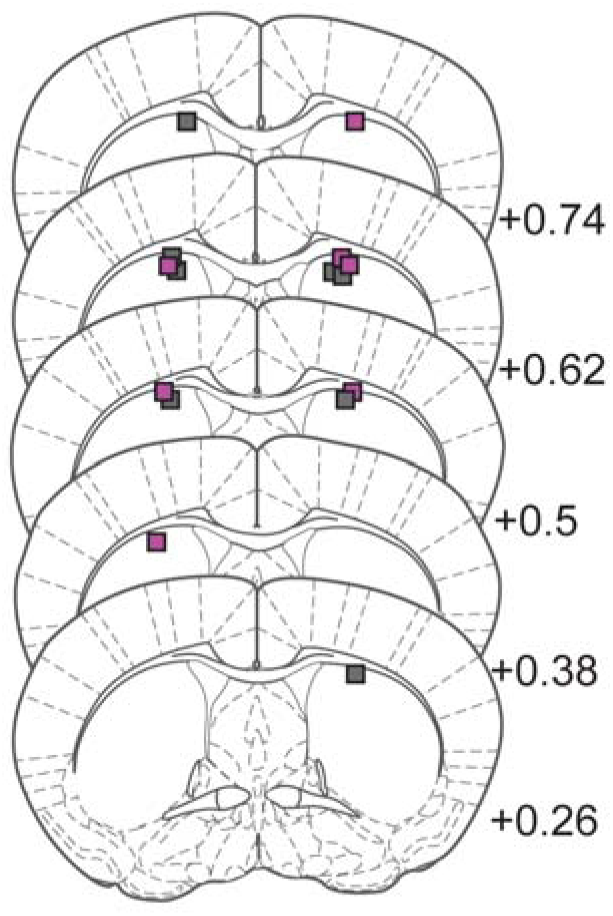
Histological analysis of optic fiber placements in nigrostriatal projection inhibition experiments. Each point represents the fiber tip position of mice expressing eYFP (N = 8 mice, gray squares) or eOPN3-mScarlet (N = 7 mice, magenta squares). Numbers indicate anterior – posterior position relative to Bregma.

